# Modulation of microRNA-502-3p significantly influences synaptic activity, dendritic spine density and mitochondrial morphology in the mice brain

**DOI:** 10.1101/2025.03.09.642262

**Authors:** Bhupender Sharma, Daniela Rodarte, Gunjan Goyal, Manuel Miranda, Rosa A. Perez, Laura Montes, Kavya Donepudi, Sreeja Eadha, Subodh Kumar

## Abstract

Synapse dysfunction is the root cause of Alzheimer’s disease (AD). Uninterrupted and regulated synapse action is crucial to maintain healthy brain function. Our previous study discovered microRNA-502-3p (miR-502-3p), a synapse-specific miRNA, highly expressed at the AD synapses. Further, *in vitro* studies unveiled the biological relevance of miR-502-3p in modulating GABA receptor function, synaptic activity and mitochondrial morphology. Current study focuses to investigate the role of miR-502-3p *in vivo* using stereotaxic injection of miR-502-3p overexpression (OE) and suppression (sponge) lentivirus (LV) into the hippocampus of C57BL/6 wild-type (WT) mice. MiR-502-3p OE and sponge LV were characterized by transducing HT22 cells followed by QRT-PCR and miRNAScope analysis of miR-502-3p. MiR-502-3p OE LV showed a very high-fold upregulation and sponge LV showed significant reduction in miR-502-3p levels. MiR-502-3p OE and sponge LV were injected into three months old WT mice brain hippocampus. Overexpression and suppression effects of miR-502-3p were studied on synaptic proteins, synapse number, mitochondrial morphology and dendritic spine density at eight-weeks post-injection. Mice injected with miR-502-3p OE LV showed reduced levels of synaptic proteins, diminished synapse formation, defective mitochondrial morphology and reduced dendritic spine density relative to control LV treated mice. While mice treated with sponge LV showed elevated levels of synaptic proteins, augmented synapses, improved mitochondrial morphology and elongated dendrites and spine density. Our *in vivo* study unveiled translational abilities of miR-502-3p to restore synapse dysfunction in AD and other neurological disorders.

## INTRODUCTION

The brain is a complex organ in the human body, composed of billions of neurons that communicate with each other through synapses. Neurons are the fundamental functional units of the brain, responsible for processing and transmitting information throughout the body. The integrity and proper functioning of synapses are essential for various cognitive processes, including learning, memory, and behavioral regulation. Synaptic dysfunction, which refers to the impairment of normal synaptic transmission and plasticity, is recognized as a key contributor to several neurodegenerative diseases, most notably Alzheimer’s disease (AD) (1, 2). Synaptic dysfunction can result from various factors including genetic mutations, environmental factors, age and microRNAs (miRNAs) (3).

MiRNAs, a class of small non-coding RNAs, have emerged as crucial regulatory molecules in modulating synaptic function (4). These small RNAs can post-transcriptionally regulate gene expression, influencing protein synthesis and synaptic plasticity. The regulation of miRNAs at synapses is critical for maintaining neuronal health and function (4, 5, 6). Disruptions in miRNA expression have been linked to alterations in synaptic activity and the progression of neurodegenerative diseases, where impaired synaptic communication contributes to cognitive decline and memory deficits (4, 7, 8). Synaptic dysfunction is one of the earliest and most prominent features of AD (1, 2). The accumulation of amyloid-beta plaques (9, 10) and tau tangles (11, 12) in the brain is associated with a progressive loss of synapses, leading to deficits in cognitive functions. Emerging evidence suggests that alterations in miRNA regulation might exacerbate synaptic dysfunction in AD, thereby accelerating neurodegenerative processes.

Our lab was the first to highlight the role of miR-502-3p in AD synapses. In previous work, we found increased miR-502-3p expression in the synaptosomal fraction from AD post-mortem brains compared to healthy controls (6). The elevated level of miR-502-3p was positively correlated with AD Braak stages. We also observed that high level of miR-502-3p levels in AD cerebrospinal fluid (CSF) were significantly associated with amyloid plaques and neurofibrillary tangles (NFTs) densities in affected brain regions (13). Additionally, we identified that miR-502-3p targets the GABA-A-receptor α1 (GABARα1) and modulates GABAergic synapse function in mouse hippocampal neurons (14). GABAergic dysfunction is well known in AD and contributes to disrupted neurological function (15). GABA, which reduces action potentials, helps maintain the balance between neural excitation and inhibition, essential for precise communication (16, 17). GABA-A-receptors, composed of pentameric subunits (α1–6, β1–3, γ1–3, δ, ρ1–3, plus minor subunits) (3, 18). The α subunit of GABA is the ligand binding site and crucial for the proper functioning of GABA neurons. GABA dysfunction is well-documented in AD, and our prior studies have shown reduced GABARα1 and VGLUT1 proteins in AD synaptosomes (14).

MiR-502-3p have a potential role in AD by modulating GABAergic and neuronal signaling pathways (3, 6, 14). The hippocampus is central to cognitive processes (19, 20), and its dysfunction contributes to AD and other dementias (21). Our previous study found high expression levels of miR-502-3p and reduced expression levels of GABARα1 in AD synaptosomes relative to controls (6). *In-silico* analysis revealed that miR-502-3p is involved in GABAergic synapse function in AD and neurodegenerative diseases (14). It targets multiple sites on GABARα1 mRNA, with gene ontology enrichment showing its significant role in nervous system development and GABAergic synapse function. Overexpression of miR-502-3p reduces GABARα1 and also decreased neuronal cell viability and increased necrotic cells. Whole-cell patch-clamp analysis of human GABA-A-receptor cells showed reduced GABA current with miR-502- 3p overexpression, indicating a negative correlation with GABAergic synapse function. Furthermore, overexpression of miR-502-3p increased AD-related protein levels, while suppression of miR-502-3p reduced AD protein and enhances GABA current (14).

Building on our previous findings (6, 13, 14, 22), we are now interested in exploring the role of miR-502-3p *in vivo* using (C57BL/6) wild type mice. To modulate miR-502-3p expression, we performed targeted delivery of miR-502-3p overexpression and suppression lentivirus into the mouse hippocampus, particularly in the CA1 area, as it is closely associated with memory and learning processes. Study purpose is to investigate the effects of miR-502-3p on neuronal activity, morphology, and molecular signaling in the mice brain. Our study unveiled protective and deleterious impacts of miR-502-3p on synaptic activity, synapse number, mitochondrial morphology and dendritic spine density that further advance our understanding on miR-502-3p’s role in synaptic function and its potential therapeutic relevance in AD.

## MATERIALS AND METHODS

### MiR-502-3p lentiviral constructs

MiR-502-3p overexpression lentivirus (miR-502-3p OE-LV) (Vector ID: VB231005-1051gew), miR-502-3p suppression sponge lentivirus (miR-502-3p sponge LV) (VB231004-1661ewa) and a scrambled control lentivirus (SC LV) (VB010000- 0009mxc) vectors were purchased from VectorBuilder, USA. All LVs were aliquoted and stored at −80^°^C until use. The miR-502-3p expression and inhibition lentiviral constructs were designed with fluorescent markers and inducible promoters to enable stable and controlled expression of miR-502-3p. Detailed information about these constructs can be found in the supplementary materials (Supplementary Information File 1). The use of inducible promoters ensures long-term stability, consistent effects on cellular processes, gene expression, and phenotype. After 24 hours post-transfection, complete DMEM media was added, and cells were incubated for an additional 48 hours at 37^°^C in a humidified atmosphere with 5% CO_2_. Transduction efficiency was assessed 48 hours post-transduction by evaluating GFP expression using fluorescence microscopy (AMG Evos Fl Microscope, E1512-155E-088). GFP-positive cells were selected to determine transfection efficiency.

### Cell culture and miR-502-3p LVs transduction

Mouse hippocampal neuronal cells (HT22) were maintained in our laboratory. Cells were cultured in Dulbecco’s Modified Eagle Medium (DMEM, Thermo Fisher Scientific) supplemented with 10% fetal bovine serum (FBS, Gibco) and 1% penicillin/streptomycin (complete media), at 37^°^C in a humidified atmosphere with 5% CO_2_. For transduction, HT22 cells (0.1 × 10^5^ cells per well) were seeded onto coverslips in a 24-well plate with DMEM containing 5% FBS and no antibiotics. After 6 hours, once the cells adhered, they were transduced with lentiviral particles of multiplicity of infection (MOI) 12. The following lentiviral constructs were used in equal quantity: (i) SC-LV (pLV[shRNA]-EGFP:T2A:Puro-U6>Scramble_shRNA), (ii) miR-502-3p OE LV (pLV[shRNA]-EGFP-U6>{hsa-microRNA-502-3p}) for miR-502-3p overexpression with GFP expression, and (iii) MiR-502-3p sponge LV (pLV[Exp]-SYN1>EGFP:{miRNA-502- 3p sponge}) designed to inhibit hsa-miR-502-3p by providing multiple binding sites for the miRNA, also with GFP expression.

### MicroRNA *in-situ* hybridization

To assess miR-502-3p overexpression in transduced HT22 cells, *in-situ* hybridization was conducted using RNAScope™ (Advanced Cell Diagnostics, Newark, CA, USA) following our previous lab publication (14). HT22 cells were seeded on poly-*L*-lysine-coated glass coverslips at a density of approximately 0.1 × 10^5^ cells per well and transduced with the scramble control, miR-502-3p overexpression, and miR-502-3p sponge lentiviruses using polybrene. Forty-two hours post-transduction, cells were fixed with 10% neutral-buffered formalin, permeabilized with PBS containing 0.1% Tween 20, and hybridized with target probes specific to hsa-miR-502-3p, mmu-U6. Signal amplification was carried out according to the manufacturer’s instructions (Advanced Cell Diagnostics, Newark, CA, USA). After hybridization, the cells were washed five times with TBST buffer at 10-minute intervals. The samples were stained with 50% hematoxylin, mounted with VectaMount mounting medium (Vector Laboratories, Newark, CA, USA), and examined using a fluorescent motorized upright microscope (Leica Microsystems).

### Quantitative reverse transcription PCR (qRT-PCR) analysis

To quantify miR-502-3p expression in LV-transduced HT22 cells and LV-injected mice, qRT-PCR was performed (14). Total RNA was extracted using TriZol reagent (Invitrogen, USA) and quantified with a Nanodrop spectrophotometer. For miRNA analysis, 2 μg of RNA was polyadenylated and reverse transcribed into cDNA using the miRNA first-strand synthesis kit (Agilent), following the manufacturer’s protocol. The qRT-PCR reaction mixture consisted of 1 μL of miR-502-3p-specific forward primer (10 μM; IDT), 1 μL of universal reverse primer (3.125 μM; Agilent), 5 μL of 2X SYBR Green PCR Master Mix (KAPA SYBR, Roche), and 1 μL of cDNA. RNase-free water was added to achieve a final volume of 10 μL. U6 small nuclear RNA (snRNA) expression was measured as an internal control to normalize miRNA expression. Each sample was prepared in triplicate and analyzed using the LightCycler 96 Real-Time PCR system (Roche Diagnostics, Indianapolis, IN, USA). The qRT-PCR cycling conditions included an initial denaturation at 95^°^C for 10 minutes, followed by 50 cycles of 95^°^C for 10 seconds, 60^°^C for 20 seconds, and 72^°^C for 20 seconds. Relative miRNA expression was calculated using the formula (2^–ΔΔCt) (14).

### Animal model

We use three months old C57BL/6 wild-type mice (total animal 21; 9 males; 12 females) for this study. All experimental procedures were approved by the Institutional Animal Care and Use Committee (IACUC) of Texas Tech University Health Sciences Center (TTUSHC) El Paso, and were in compliance with the NIH guidelines. Mice were housed under standard laboratory conditions with a 12-hour light/dark cycle and provided access to food and water at Laboratory Animal Resource Center at TTUHSC El Paso.

### Stereotaxic surgery

Mice were divided into three groups, each consisting seven mice. The mice groups were injected with following lentivirus: Group 1. Scramble control LV (SC LV), Group 2. miR-502-3p OE LV, and Group 3. miR-502-3p sponge LV. The three-months old mice were anesthetized with isoflurane (0-4% induction, 1-2% maintenance). The mice were carefully placed on the nose cone, where isoflurane (2%) was administered through the SomnoSuite. Eye lubricant was applied to protect their eyes from dryness during the procedure. A weight-based subcutaneous dose of analgesic flunixin meglumine (2.5 mg/kg) was administered prior to surgery to minimize postoperative pain. Their fur was trimmed with a trimmer in preparation for surgery. The mice were then placed on a stereotaxic apparatus (Stoelting Co., Wood Dale, IL) for precise localization of the hippocampal CA1 region. The ear bars were used to securely stabilize the mouse head on the stereotaxic instrument, ensuring proper alignment and stability for precise targeting. Anesthesia was maintained with isoflurane (0.9-3%). The surgical site three times with chlorhexidine and alcohol. A small incision was made along the midline of the skull and hydrogen peroxide (3% v/v) was applied to visualize the bregma and lambda. The skull was then carefully adjusted for the procedure. Subsequently, a 2 mm hole was drilled using a micromotor drill (Foredom MH-130) at the coordinates: anterior-posterior (AP): −2.03 mm, medial-lateral (ML): −1.7 mm and dorsal-ventral (DV): −1.3 mm, based on the Paxinos and Franklin Atlas for precise targeting of the hippocampal CA1 region (23). The 2 μL of either SC LV, miR-502-3p OE LV and miR-502-3p sponge LV were injected into the hippocampal CA1 region on both sides using a 5 μL, Hamilton syringe (Hamilton, Reno, NV). The LVs were injected at a rate of 150 nL/ min over a period of 5 minutes to minimize tissue damage and avoiding LVs leakage. After the injection, the incision was closed with sterile sutures, and a triple antibiotic ointment was applied to prevent infection. The mice were allowed to recover on a heating pad and were closely monitored for any postoperative complications.

### Confirmation of miR-502-3p LV injections

Five mice from each group (total 15 wild-type C57BL/6 mice; 6 males; 9 females) were euthanized two-months after LV-injection using CO_2,_ followed by cervical dislocation. After euthanasia, the brains were carefully removed and processed for further analysis. Two LV-injected mice brains from each group (total 6; 3 males; 3 females) were cryoprotected in a 30% sucrose solution at 4^°^C. The cryoprotected frozen brains were sectioned coronally at 20 μm using a microtome (Leica, CM1850). Sections containing the hippocampal CA1 region were collected and stored in cryoprotectant solution at −20^°^C until further analysis, including green fluorescent protein (GFP) and immunostaining. Some sections were mounted on glass slides and examined for GFP expression to confirm LV-injection. Whole-brain images were captured using an APX100-HCU (Olympus, Evident Corporation, Nagano, Japan) with a 40X objective to confirm LVs injection. The middle region of the mouse brain, containing the injection sites, was isolated from remaining three LV-injected wild-type mice from each group. Hippocampal and cortical tissues were separated from these brains and stored at −20^°^C for RNA (qRT-PCR) and protein (western blotting) analysis. Total RNA was extracted using the Trizol method. RNA concentration and integrity were assessed using a NanoDrop spectrophotometer (Thermo Fisher Scientific). Reverse transcription was performed using the High-capacity cDNA reverse transcription kit (Thermo Fisher Scientific) and the miRNA First strand synthesis kit (Agilent). To analyze the expression of miR-502-3p by qRT-PCR, gene-specific primers for miR-502-3p and U6 (a housekeeping gene), were commercially synthesized by Integrated DNA Technologies (IDT), Coralville, IA, USA (Supplementary Information File 2). Reactions were performed using the Kappa SYBR master mix (Roche) on a Real-Time PCR system (Roche). The fold change in gene expression was calculated using the ΔΔCt method.

### Synaptic protein study- Immunoblotting analysis

For immunoblotting analysis, mouse brain hippocampal tissues stored at −20^°^C from each group were used and suspended in RIPA buffer (Thermo Scientific) supplemented with 1X protease inhibitor and EDTA. The samples were sonicated on ice (Amplitude 50%, Pulse 5 sec on/off) for 10 seconds. A total of 30 μg of protein samples were mixed with 4X LDS sample loading buffer (Novex) in a 4:1 (v/v) ratio and subjected to SDS-PAGE. Electrophoresis was performed using a mini protein tetra cell (Bio-Rad) at 90 V for 2 hours. A transfer sandwich was prepared with the gel facing the anode and a PVDF membrane facing the cathode. Protein transfer was carried out for 2 hours at 80 V at 4^°^C in transfer buffer using the Mini Protein Tetra Cell (Bio-Rad) transfer system. Subsequently, the PVDF membrane was blocked with 5% (w/v) bovine serum albumin (BSA) in Tris-buffered saline with 0.1% Tween 20 (1X TBS-T). The blots were incubated overnight at 4^°^C with anti-mouse primary antibodies, including GABARα1 (1:500), Gephyrin (1:3000), PSD95 (1:1000) and Synaptophysin (1:500), prepared in 5% BSA (w/v; 10 mL). Detailed antibody information and dilution factors are provided (Supplementary Information File 2). Following primary antibody incubation, the blots were washed five times with TBS-T and incubated for 2 hours at room temperature with secondary antibodies, anti-mouse/anti-rabbit IgG-peroxidase (Sigma; 1:10,000) prepared in 5% BSA (w/v). After washing the blots five times with TBS-T, protein bands labeled with secondary antibodies were detected using Clarity™ enhanced chemiluminescence (ECL; Bio-Rad, USA) western blotting substrate. The blots were then visualized by exposure to dark conditions in a luminescent image analyzer (Amersham Imager 680; GE Healthcare Bio-Sciences, Sweden). Protein band intensities were quantified using ImageJ software (1.54d, Java 1.8.0_345; http://imagej.org) for densitometry analysis, as described in more detail in our previous study. Relative protein expression levels were normalized to β-tubulin as a loading control.

### Dendritic spine density- Golgi-cox staining

Two wild-type mice from each LV-injected group, (total of 6 wild-type mice; 3 males and 3 females, C57BL) were anesthetized with isoflurane (0-4% for induction, 2% for maintenance via nose cone) and trans-cardially perfused with normal saline to clear the blood from the vasculature, followed by 4% paraformaldehyde (PFA) in 0.1M phosphate buffer (pH 7.4) for 4 minutes (Welch 3200). After perfusion, the brain was carefully removed. Tissue samples were collected from one LV-injected mouse from each group, fixed in 4% paraformaldehyde at 4^°^C overnight, and stored for transmission electron microscopy (TEM) analysis. Additionally, LV-injected one mice brain from each group was processed for Golgi-cox staining. A tissue block (∼5 mm thick) from the middle region of the mouse brain, containing injection sites, was collected. The mice brain tissue was rinsed quickly with milli-Q water to remove blood from the surface. The tissues for Golgi-cox staining were then immerged in impregnation solution A + B (FD Rapid Golgi Stain Kit, FD NeuroTechnologies, Inc., Columbia, MD). The impregnation solution was replaced after 24 hours, and the tissue was stored at room temperature in the dark for 2 weeks. The container was gently swirled twice a week for 30 seconds to ensure uniform impregnation. After 2 weeks, the tissues were transferred into solution C. Solution C was replaced after 24 hours, and the tissue was stored in the dark at room temperature for 7 additional days. The samples were then sent to FD NeuroTechnologies, Inc. for sectioning, mounting, and staining. Tissue sections (100 µm) containing the hippocampal CA1 region were used to analyze the effects of LV- injection on neurons. The entire mouse brain (4X), hippocampal injection site (10X), and dendritic spine density and morphology were quantified using an APX100-HCU (Olympus, Evident Corporation, Nagano, Japan) with a 40X objective.

### Synapse number and mitochondrial morphology- Transmission electron microscopy analysis

To assess the impact of miR-502-3p on mitochondrial number and size, mitochondria were analyzed in hippocampal tissue near the LV injection site for TEM analysis. Paraformaldehyde (4%) fixed LV-injected mouse brain (C57BL; 3 males, as described above in Section 2.4) hippocampal tissue one from each group were used for mitochondrial analysis. The hippocampus and the cerebral cortex were isolated and cut into ∼1 mm³ cubes from 4% paraformaldehyde fixed mouse brains. The tissues were again fixed in a solution containing 2% paraformaldehyde and 2.5% glutaraldehyde in 0.05 M cacodylate buffer, followed by post-fixation with 1% osmium tetroxide, and then embedded in LX-112 resin. Ultrathin sections were cut, stained with uranyl acetate and lead citrate, and examined using a Hitachi H-7650 TEM at 60 kV, located at the College of Arts and Sciences Microscopy Facility, Texas Tech University, Lubbock, Texas, USA. TEM images were used to quantify synapse numbers in the cortex and hippocampal regions of the following groups: Scramble control (SC) LV-injected cortex; Scramble control LV-injected hippocampus; miR-502-3p overexpression (OE) LV-injected cortex; miR-502-3p OE LV-injected hippocampus; miR-502-3p sponge LV-injected cortex; miR- 502-3p sponge LV-injected hippocampus. Images at a 500 nm scale were selected for quantification. Synapse counts focused on the organization of synapses (including mitochondria, synaptic vesicles, and synaptic clefts), and the average number of synapses per slide was determined for the Sc, miR-502-3p OE, and miR-502-3p sponge LV-injected cortex and hippocampus.

### Statistical Analysis

All data are presented as mean ± standard error of the mean (SEM). Statistical analyses were performed using GraphPad Prism (GraphPad Software, Inc., La Jolla, CA). For comparison between SC LV, miR-502-3p OE LV and miR-502-3p sponge LV groups, one-way analysis of variance (ANOVA) followed by Tukey’s post hoc test was used. Data was also compared between SC LV versus miR-502-3p OE LV, SC LV versus miR-502-3p sponge LV and miR-502-3p OE LV versus miR-502-3p sponge LV using student t-test. All parameters A p-value of < 0.05 was considered statistically significant.

## RESULTS

### Characterization of miR-502-3p overexpression and suppression LVs

The genetic map of the lentiviral vectors (LVs) highlighting key components, including the miR-502-3p precursor sequence, promoter regions, and the GFP reporter gene, which facilitated monitoring of successful transduction. This construct was used to transduce HT22 cells, resulting in the observed upregulation of miR-502-3p expression, as confirmed by qRT-PCR analysis. Following LV transduction to HT22 cells, robust GFP expression was observed, indicating successful transduction (Figure 1). Figure 1.A showed the schematic map of each LV’s: scramble control LV, miR-502-3p OE LV and miR-502-3p sponge LV. The cells displayed bright green fluorescence protein under a fluorescence microscope, confirming effective LV-mediated gene transfer (Figure 1.B). Additionally, the overexpression of miR-502-3p was confirmed using miRNA *in-situ* hybridization (miRNAscope). The hybridization probe specific to miR-502-3p revealed strong signals within the HT22 cells transduced with miR-502-3p LVs, which were predominantly localized in the cytoplasm (Figure 1.C). This indicates that miR-502-3p was effectively overexpressed in the cells, and its expression pattern was detectable using the *in-situ* hybridization technique. Quantitative PCR analysis revealed a significant increase in miR-502-3p expression (P < 0.001) in HT22 cells transduced with the miR-502-3p overexpression LVs (Figure 1.D). The miR-502-3p levels in the experimental group were markedly higher compared to the control cells, demonstrating successful overexpression. These findings confirm the effective transduction and overexpression of miR-502-3p in HT22 cells, as evidenced by the substantial upregulation detected by qRT-PCR. Collectively, these results provide a reliable basis for further investigation into the functional impact of miR-502-3p in mice.

**Figure 1:**
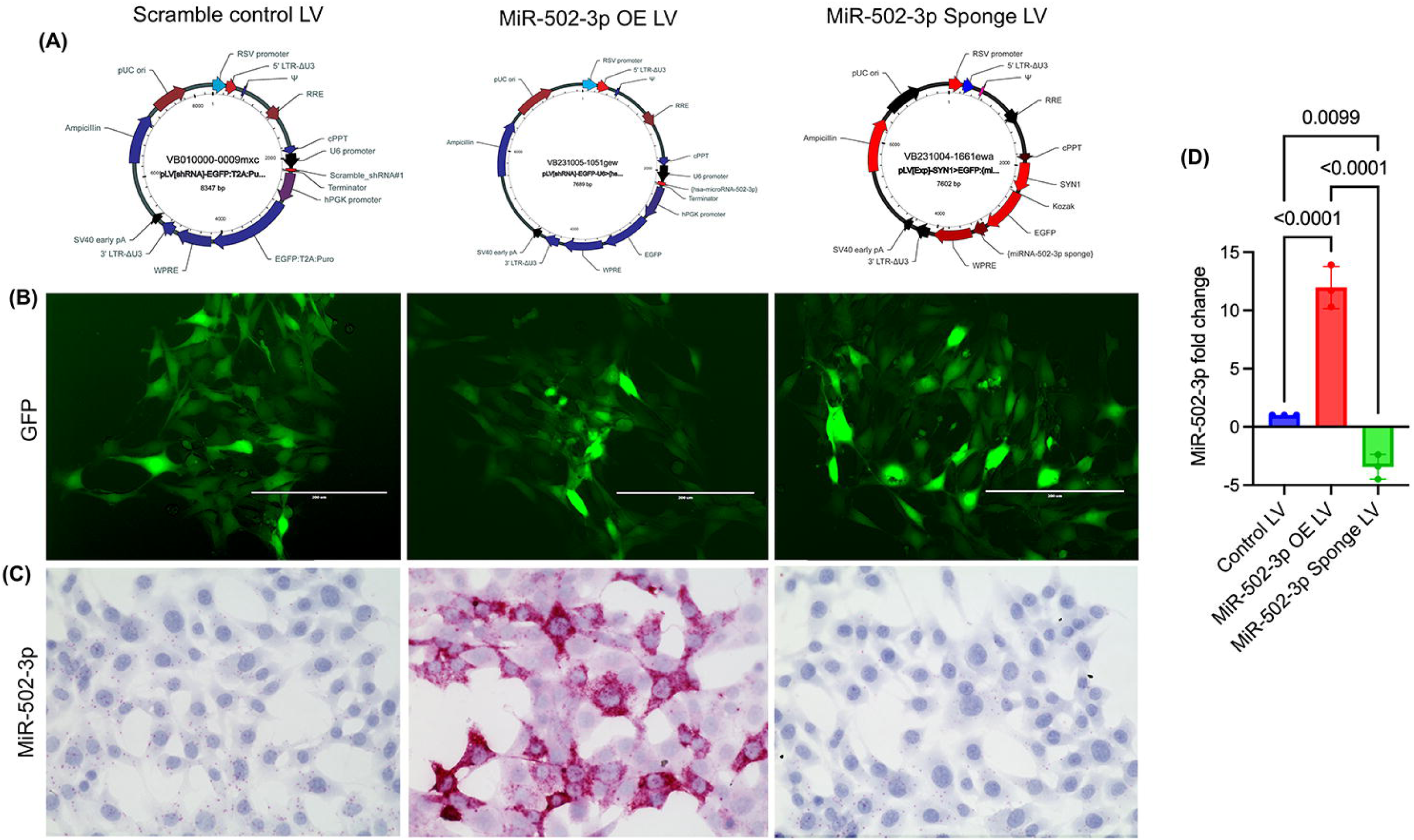
Confirmation of miR-502-3p overexpression after lentiviral vector transduction. (A) Genetic map of the lentiviral vector (LV) used for transduction of HT22 cells, highlighting the scrambled control lentivirus (SC LV), miR-502-3p overexpression lentivirus (miR-502-3p OE LV) and miR-502-3p suppression lentivirus (miR-502-3p Sponge LV), promoter regions, and GFP reporter gene for monitoring transduction efficiency. (B) HT22 cells transduced with miR-502-3p LVs displayed GFP expression, bright green fluorescence under a fluorescence microscope, confirming successful transduction. (C) *In-situ* hybridization (miRNA scope) was used to assess miR-502-3p overexpression. The miR-502-3p-specific probe showed strong signals in the cytoplasm of the transduced HT22 cells, confirming effective miRNA overexpression (scale bar = 200 µm). (D) Quantitative PCR analysis of miR-502-3p expression in HT22 cells transduced with the miR-502-3p overexpression LVs showed a significant increase in miR-502-3p levels compared to control cells (**P < 0.001), confirming successful overexpression. These results demonstrate effective transduction and upregulation of miR-502-3p in HT22 cells, providing a foundation for future functional studies of miR-502-3p.

### Detection of miR-502-3p overexpression and suppression LV in mice hippocampus

Stereotaxic injections were performed in three-month-old wild-type C57BL/6 mice (21 total; 9 males, 12 females), with scrambled control lentivirus (SC LV), miR-502-3p overexpression (miR-502-3p OE LV) and miR-502-3p suppression LV (miR-502-3p sponge LV) injected into the hippocampal CA1 region. Two months post-injection, the effectiveness of lentiviral delivery was assessed by analyzing viral transduction efficiency and the spatial distribution of lentiviral particles. The overall experimental design for the LV injection is shown (Figure 2.A). Sections containing the hippocampal CA1 region were examined for GFP expression to confirm the success of the LV injection. The hippocampal CA1 injection site was marked with white squares for clarity. Fluorescent microscopy of brain sections revealed widespread GFP expression across all experimental groups, with strong fluorescence in the CA1 region and extending into adjacent areas (Figure 2.B). The GFP marker confirmed successful lentiviral delivery in all groups, and histological analysis of brain sections verified the precise localization of the injection site. Whole-brain images were captured using an APX100-HCU microscope (Olympus, Evident Corporation, Nagano, Japan) at 40X magnification (Figure 2.B). Total RNA was then extracted from hippocampal tissue near the injection sites in three mice from each LV-injected group (total 9 mice; 3 males, 6 females). The results showed a significant 3.3-fold overexpression of miR-502-3p (P = 0.0119) in the hippocampus of mice injected with miR-502-3p OE LV, whereas miR-502-3p expression was reduced to 0.5-fold (P = 0.0117) in the hippocampal tissue of mice injected with miR-502-3p sponge LV, compared to the Scramble LV-injected group (Figure 2.C).

**Figure 2:**
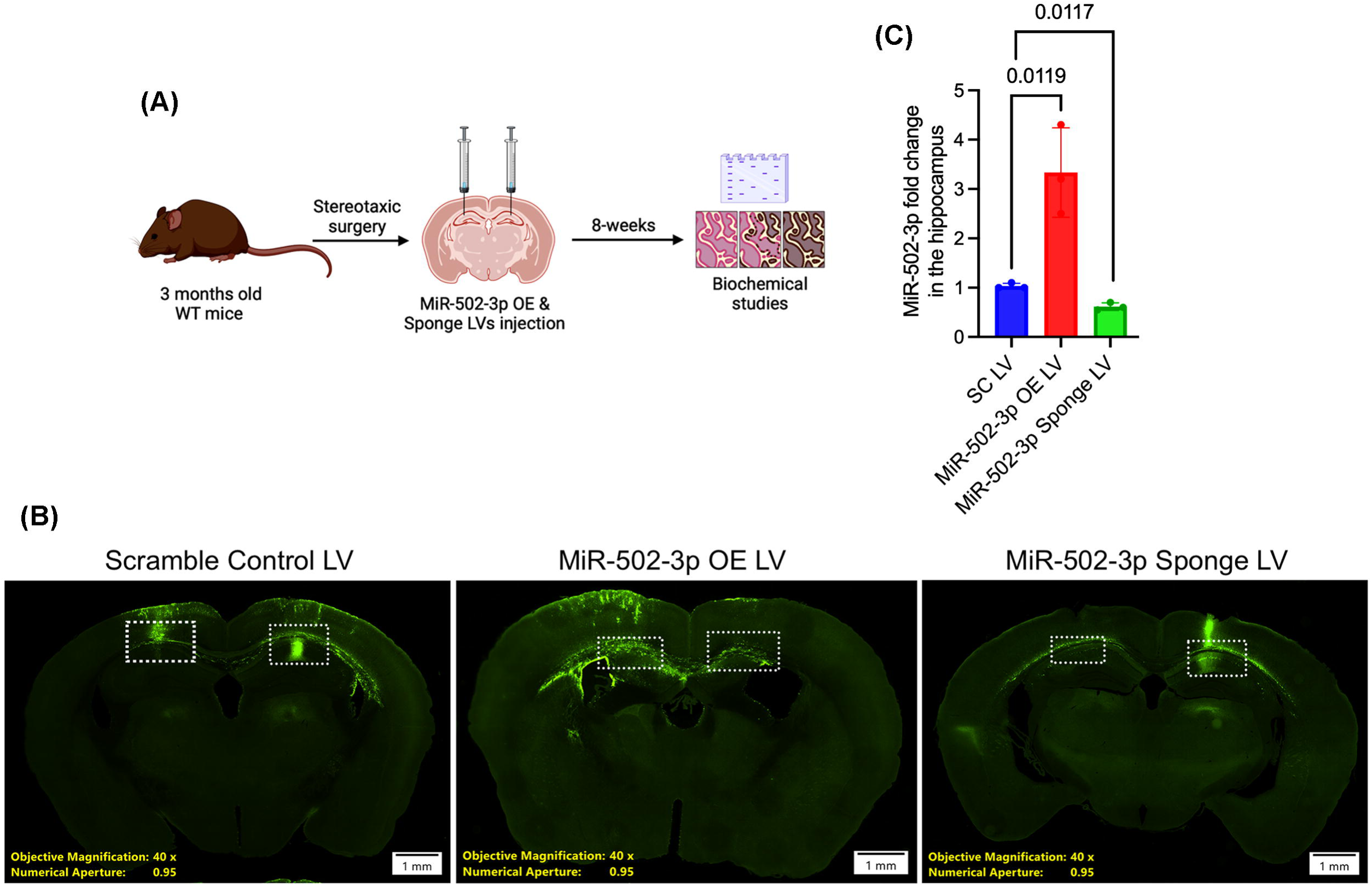
Lentiviral injection in the mice hippocampal CA1 region. (A) Schematic representation of the experimental design for stereotaxic injections of LV into the hippocampal CA1 region of two-month-old wild-type C57BL mice. Mice were injected with either scrambled control lentivirus (Scramble LV), miR-502-3p overexpression lentivirus (miR-502-3p-OE LV) or miR-502-3p sponge lentivirus (miR-502-3p sponge LV). Two months post-injection, the effectiveness of lentiviral delivery was assessed by analyzing GFP expression and miR-502-3p levels. (B) Representative fluorescent images of brain sections from wild-type mice injected with miR-502-3p-OE LV, miR-502- 3p sponge LV, or Scramble LV, showing widespread GFP expression in the CA1 region (marked with white squares) and surrounding areas, confirming successful lentiviral transduction. Images were captured using a 40X objective (scale bar = 1 mm). (C) Quantitative PCR analysis of miR-502-3p expression in hippocampal tissue from LV-injected mice. miR-502-3p levels were significantly elevated by 3.3-fold (P = 0.0119) in mice injected with miR-502-3p-OE LV and significantly reduced to 0.5-fold (P = 0.0117) in mice injected with miR-502-3p sponge LV, compared to the Scramble LV control group. These results confirm the successful delivery and functional expression of miR- 502-3p in the hippocampus following LV injection.

### Impact of miR-502-3p overexpression and suppression on synaptic proteins

Western blot analysis was conducted to confirm changes in synaptic protein expression across the miR-502-3p OE LV, miR-502-3p sponge LV, and scramble LV groups. Significant reductions in the protein levels of GABRα1, Gephyrin, synaptophysin and PSD-95 were observed in the miR-502-3p OE LV group compared to the controls (Figure 3.A). These findings suggest that miR-502-3p plays a role in regulating synaptic protein expression, specifically affecting synaptic integrity and plasticity. In contrast, the miR-502-3p sponge LV group exhibited significantly increased expression of all proteins: GABRα1, Gephyrin, PSD95 and synaptophysin supporting the hypothesis that miR-502-3p regulates synaptic protein levels (p < 0.05). This increased protein expression in the sponge group further highlights the potential of miR-502-3p as a key regulator of synaptic function and plasticity. Densitometry analysis of protein bands using ImageJ software showed that protein levels of GABRα1, Gephyrin, PSD95 and synaptophysin were significantly reduced in the miR-502-3p OE LV group, while protein expression was significantly increased in the miR-502-3p sponge LV group (Figure 3.B). Protein levels were normalized to β-tubulin to correct for variations in loading, and densitometry analysis was performed. These results underscore the functional impact of miR-502-3p on synaptic protein expression and provide further insights into its role in synaptic integrity and plasticity.

**Figure 3:**
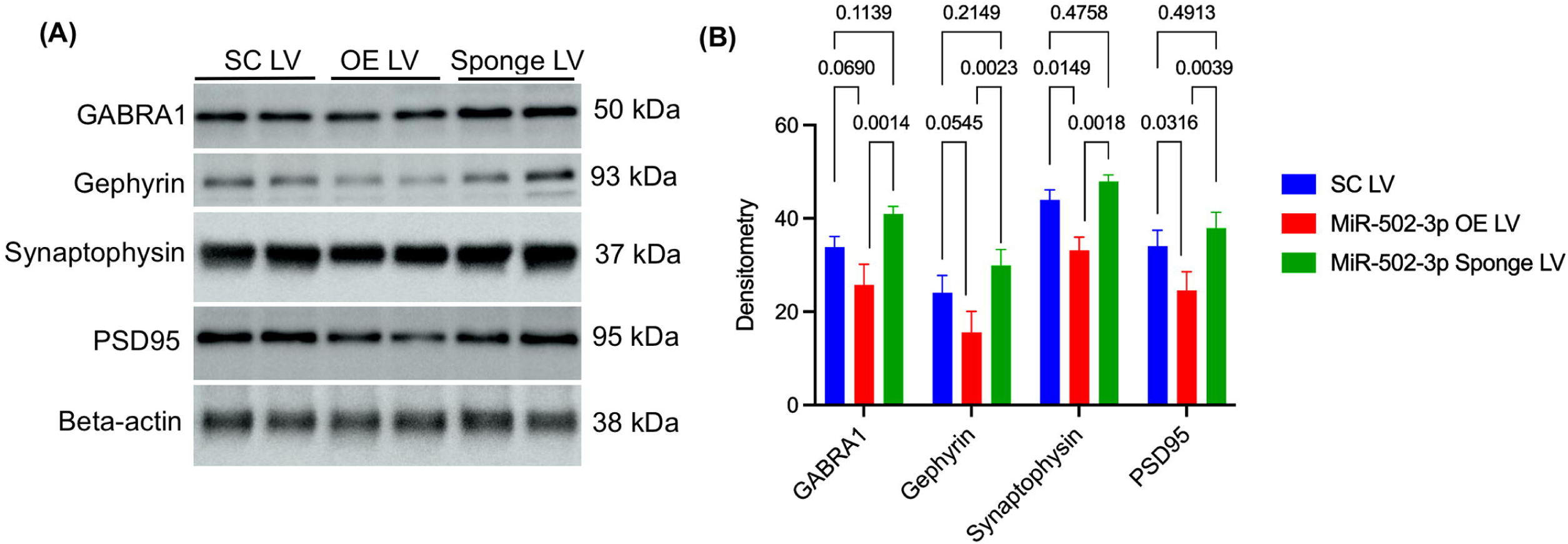
Effect of miR-502-3p overexpression on synaptic markers. (A) Western blot analysis showing protein expression of synaptic markers, including GABRα1, Gephyrin, synaptophysin and PSD-95 in the hippocampal tissue from mice injected with Scramble control LV, miR-502-3p OE LV and miR-502-3p sponge LV. A significant reduction in protein levels was observed for all markers in the miR-502-3p OE LV group compared to the control groups, indicating the potential role of miR-502-3p in regulating synaptic protein expression and synaptic plasticity. Conversely, the miR-502-3p sponge LV group exhibited significantly increased expression of these proteins, supporting the hypothesis that miR-502-3p modulates synaptic protein levels. (B) Densitometry analysis of western blot bands performed using ImageJ software, with protein levels of GABRα1, Gephyrin, synaptophysin and PSD-95 normalized with β-actin expression to account for loading differences. A significant reduction in protein expression was observed in the miR-502-3p OE LV group, while the miR-502-3p sponge LV group showed a marked increase in expression of all proteins (**P < 0.05). These findings highlight the regulatory role of miR-502-3p in synaptic protein expression and its impact on synaptic integrity and plasticity.

### Impact of miR-502-3p overexpression and suppression on dendrites morphology and spine density

To assess the impact of miR-502-3p modulation on neuronal morphology, Golgi-Cox staining was performed. Figure 4.A displays an image of the complete mouse brain at 4X magnification, offering an overall view. Overexpression of LV-miR-502-3p led to the formation of large clumps of stained neurons, which may indicate cell death, abnormal neuronal migration, or excessive damage in the hippocampus CA1 region. At 10X magnification (Figure 4.B), the injection site in the hippocampus is shown. miR- 502-3p injection caused neuronal clumping, whereas the control group exhibited a more uniform distribution of neurons. In contrast, miR-502-3p Sponge LV injection resulted in an increased and more uniform distribution of neurons in the hippocampus CA1 region. The same hippocampal region is shown at 20X magnification (Figure 4.C). In the miR- 502-3p OE LV injected group, clumped neurons were observed, while the SC LV-injected group displayed a more uniform neuronal distribution. Additionally, the miR- 502-3p sponge LV (sponge LV) injection led to increased neuronal length and enhanced network formation. At 40X magnification, dendritic complexity and spine density in CA1 pyramidal neurons were analyzed. SC LV-injected neurons exhibited a healthier network compared to those in the miR-502-3p OE LV-injected brains, where significant damage to the neuronal network was observed. In contrast, the miR-502-3p sponge LV injection showed an increase in both neuronal length and spine density (Figure 4.D). These results suggests that inhibiting miR-502-3p enhances synaptic growth and dendritic complexity compared to baseline measures, serving as a control for the experimental manipulations (Figure 4.A-D).

**Figure 4:**
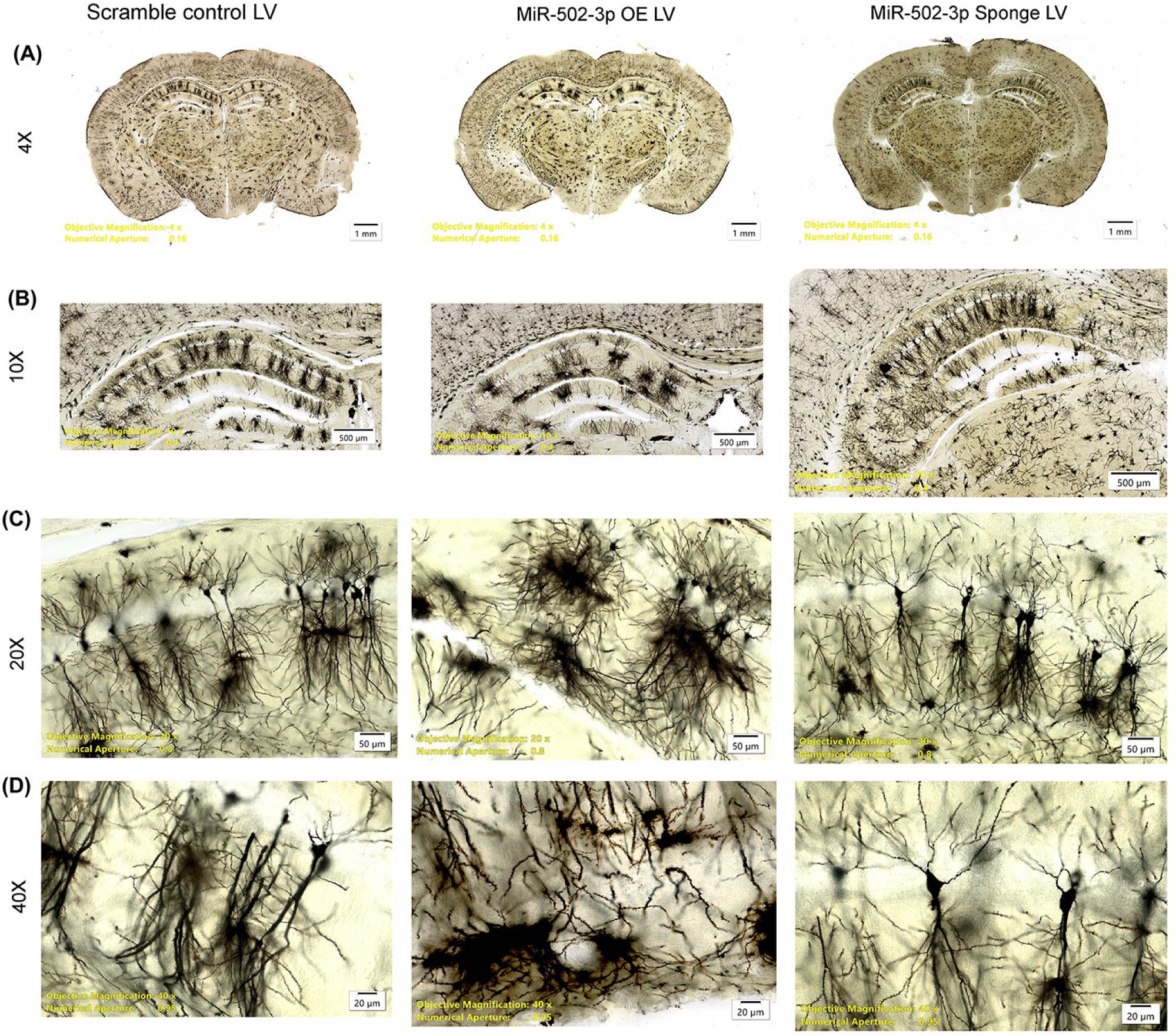
Effect of miR-502-3p overexpression on synaptic growth and dendritic complexity. (A) Low magnification image (4X) of the complete mouse brain injected with Scramble control LV, miR-502-3p OE LV and miR-502-3p sponge LV. Overexpression of miR-502-3p results in the formation of large clumps of stained neurons, which may suggest neuronal damage, cell death, or abnormal migration, particularly in the hippocampus CA1 region. (B) Intermediate magnification (10X) image focusing on the hippocampus injection site. miR-502-3p overexpression leads to clustering of neurons, while the control group displays a more uniform distribution. miR- 502-3p sponge LV injection results in an even distribution of neurons in the hippocampus CA1 region. (C) Higher magnification (20X) image showing the same hippocampal region. In the miR-502-3p overexpression group, neuronal clumping is evident, whereas the Scramble LV group shows a more uniform neuronal arrangement. The miR-502-3p sponge LV group exhibits increased neuronal length and enhanced network formation. (D) High magnification (40X) analysis of dendritic complexity and spine density in CA1 pyramidal neurons. Scramble LV-injected neurons show a healthier network compared to the miR-502-3p-OE LV group, which demonstrates significant damage. miR-502-3p sponge LV injection results in increased dendritic length and spine density. These findings suggest that inhibition of miR-502-3p promotes synaptic growth and dendritic complexity relative to baseline controls (Figure 4.A-D).

### Impact of miR-502-3p overexpression and suppression on synapse number

To examine the effects of miR-502-3p on synaptic ultrastructure, we conducted TEM analysis on hippocampal samples from SC LV, miR-502-3p OE LV and miR-502- 3p sponge LV injected mice. TEM images at 6000x (Figure 5.A) and 10000x magnification (Figure 5.B) are shown. Using eight distinct TEM fields, we counted the synapse numbers across different conditions. The miR-502-3p sponge LV group displayed an increase in synapse count (p = 0.0051). The average synapse count, where the miR-502-3p sponge LV group showed more synapses (Figure 5.C). These findings highlight the differential impact of miR-502-3p suppression on synaptic structure and function in the hippocampus.

**Figure 5.**
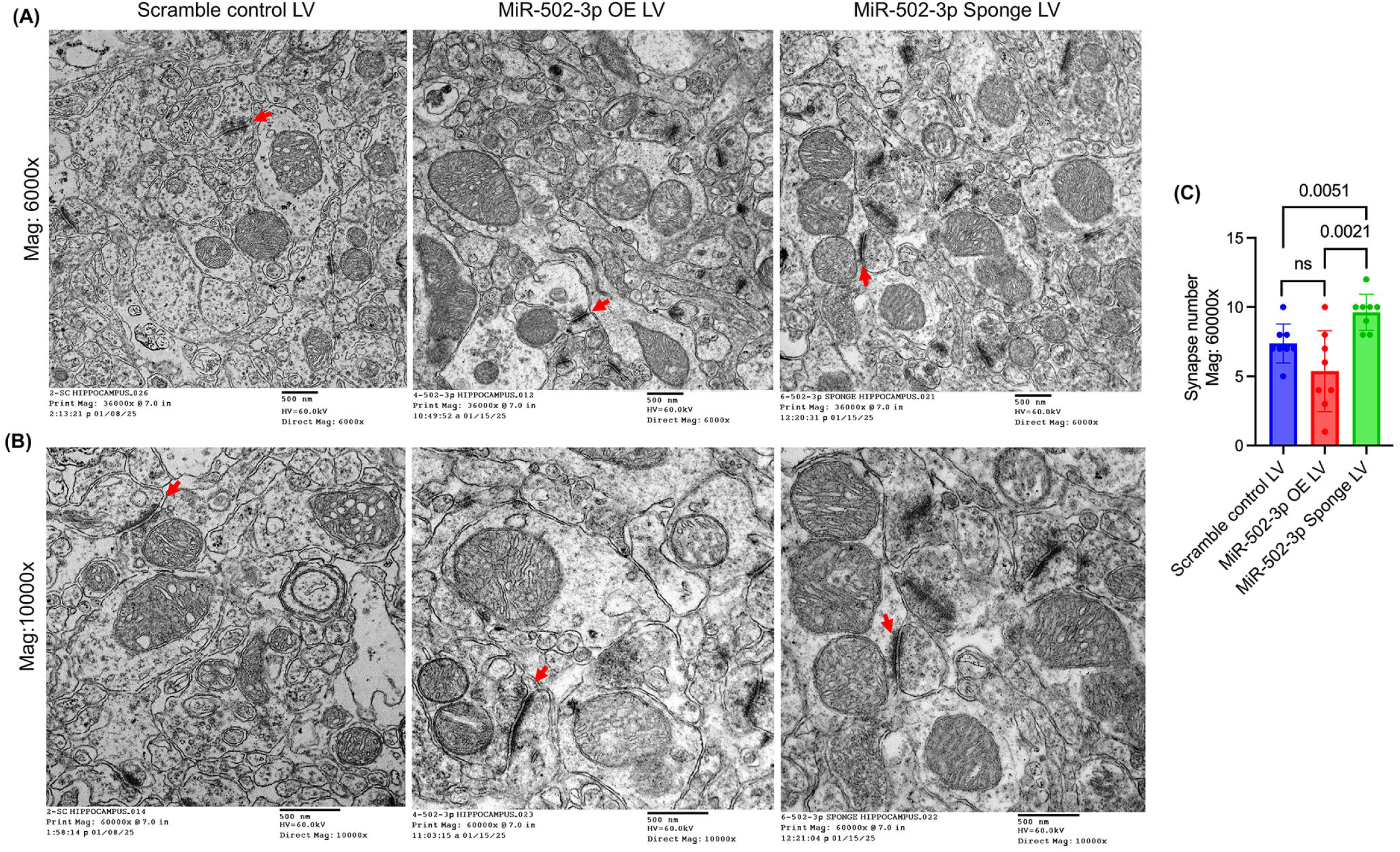
Analysis of synapse number in the hippocampal region. (A) TEM image of mice brain hippocampal region injected with Scramble control LV, miR-502-3p OE LV and miR-502-3p sponge LV at (A) 6000x magnification and (B) 10000x magnification showing synapse ultrastructure with red arrow. (C) Quantification of synapse numbers across different conditions, showing a significant increase in synapse count in the miR- 502-3p sponge LV group compared to the OE LV and scramble control LV group (p = 0.0051). These results highlight the impact of miR-502-3p suppression on synaptic structure and function in the hippocampus.

**Figure 6.**
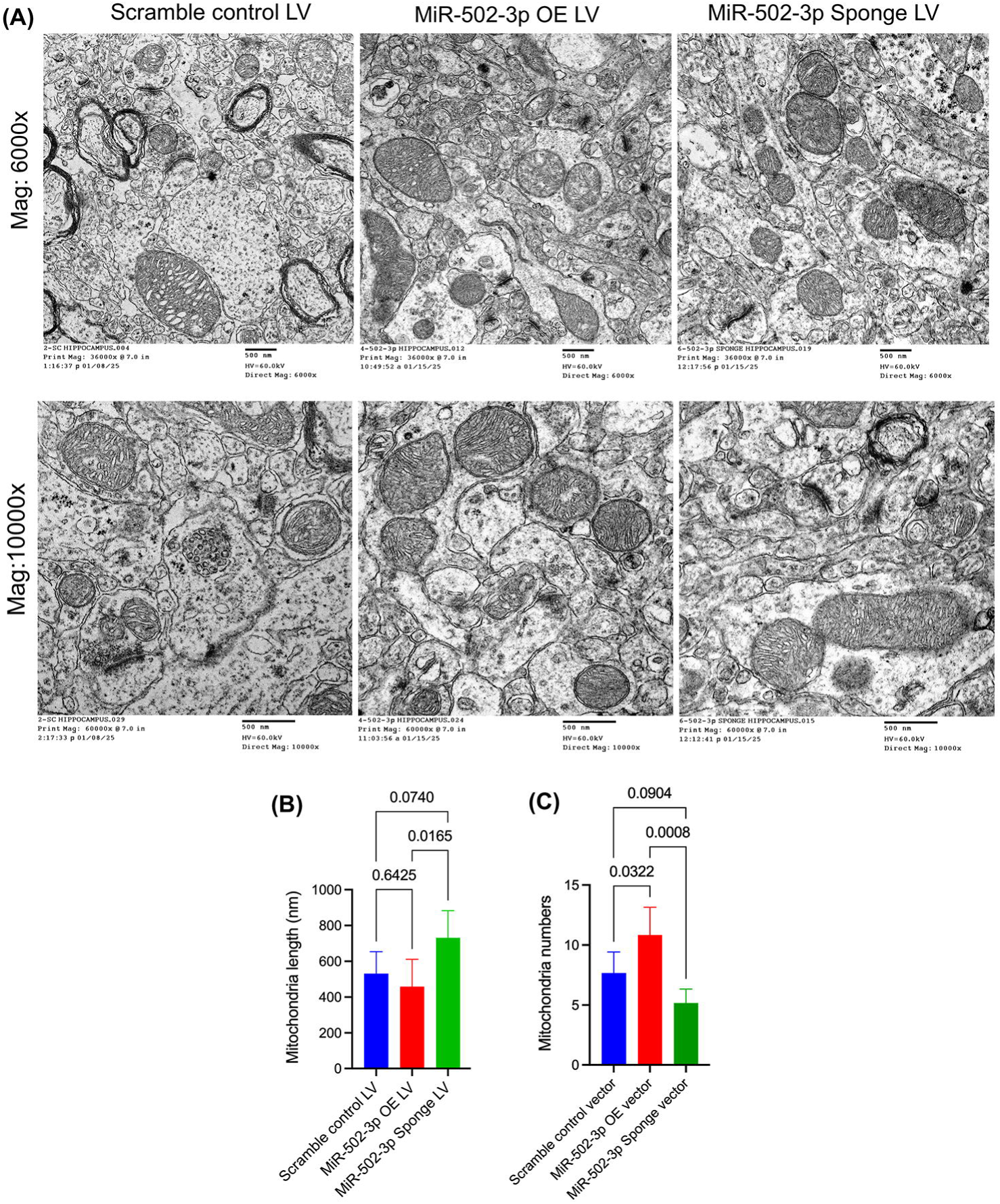
Analysis of mitochondrial morphology in the hippocampal region. (A) TEM image of mice brain hippocampal region injected with Scramble control LV, miR- 502-3p OE LV and miR-502-3p sponge LV at (A) 6000x magnification and 10000x magnification showing mitochondrial morphology. (B) Quantification of mitochondrial length across different conditions, showing a significant increase in mitochondrial length in the miR-502-3p sponge LV group compared to the scramble control and miR-502-3p OE LV group (p = 0.0165). (C) Quantification of mitochondrial number counts in all three groups, mitochondrial number was significantly high in miR-502-3p OE treated mice and reduced in Sponge LV treated mice. These results highlight the impact of miR-502-3p suppression on mitochondrial structure and function in the hippocampus.

### Impact of miR-502-3p overexpression and suppression on mitochondrial morphology

To examine the effects of miR-502-3p on mitochondrial ultrastructure, we conducted TEM analysis on hippocampal samples from SC LV, miR-502-3p OE LV and miR-502-3p sponge LV injected mice. TEM images at 6000X and 10000X magnification (Figure 5.A) are shown. Using eight distinct TEM fields, we counted the mitochondrial numbers and length across different conditions. The miR-502-3p sponge LV group displayed a significant increase in mitochondrial length (p = 0.0165) (Figure 5.B). The average mitochondria counts were significantly reduced in miR-502-3p sponge LV group while mitochondria number were more in the miR-502-3p OE mice relative to control and sponge LV treated mice (Figure 5.C). These findings highlight the differential impact of miR-502-3p suppression on mitochondrial morphology in the hippocampus. All together, these finding unveiled that overexpression of miR-502-3p displayed detrimental effects on synaptic markers while inhibition of miR-502-3p displayed protective impacts on the synaptic parameters in mice brain.

## DISCUSSION

This study explores the role of miR-502-3p in synaptic function by using LV-based overexpression and suppression models in both cultured HT22 cells and the hippocampus of wild-type mice. The results shed light on how miR-502-3p modulates synaptic integrity and neuronal morphology, providing new insights into its potential role in neurodegenerative diseases. Our results highlight the significant impact of miR-502- 3p modulation on synaptic integrity, protein expression, and ultrastructure, providing valuable insights into its potential role in synaptic plasticity and neuronal health. The successful transduction of HT22 cells with miR-502-3p overexpression lentivirus (miR- 502-3p OE LV) was confirmed through GFP expression, qRT-PCR, and *in-situ* hybridization. These results demonstrate that the lentiviral vectors effectively mediated gene transfer, leading to robust miR-502-3p overexpression *in-vitro* a result that aligns with previous studies utilizing similar LV for gene delivery (24, 25, 26). The significant increase in miR-502-3p levels in the transduced HT22 cells is consistent with previous findings from other miRNA studies, demonstrating the reliability of this approach for investigating the functional consequences of miRNA modulation (27). The significant upregulation of miR-502-3p observed in the experimental group further supports the effectiveness of this approach, providing a reliable foundation for subsequent *in-vivo* analysis. In the *in-vivo* model, we observed successful lentiviral delivery to the hippocampal CA1 region as evidenced by GFP expression, indicating efficient viral transduction across all experimental groups. The quantification of miR-502-3p expression in the hippocampal tissue confirmed significant overexpression in the miR- 502-3p OE LV group and a reduction in expression in the miR-502-3p sponge LV group, relative to the scrambled control group. These results confirm that our LV-based approach successfully altered miR-502-3p expression in the targeted hippocampal region, setting the stage for assessing its functional impact on synaptic structure and plasticity.

Western blot analysis revealed a significant downregulation of key synaptic proteins, such as GABRα1, synaptophysin, Gephyrin, and PSD-95, in the miR-502-3p OE LV group, suggesting that overexpression of miR-502-3p impairs synaptic integrity and plasticity. Conversely, the miR-502-3p sponge LV group exhibited a marked increase in the expression of these proteins, supporting the hypothesis that suppression of miR-502-3p enhances synaptic function. This is in line with previous studies indicating that miRNAs can regulate synaptic protein expression and are involved in synaptic dysfunction in neurodegenerative diseases (4, 6, 13, 14, 22). These findings highlight miR-502-3p as a potential regulator of synaptic protein expression and provide evidence for its involvement in maintaining synaptic integrity.

Golgi-Cox staining provided valuable insights into the effects of miR-502-3p modulation on dendritic morphology and spine density. The miR-502-3p OE LV group showed signs of neuronal clumping, suggesting possible cell death or abnormal neuronal development in the hippocampus. This finding is supported by previous studies showing that the overexpression of certain miRNAs can lead to neuronal stress and cell death (28). In contrast, the miR-502-3p sponge LV group displayed enhanced dendritic growth and spine density, suggesting that suppression of miR-502-3p promotes synaptic development and neuronal network formation. These observations are further supported by the increase in neuronal length and enhanced network formation in the sponge group, highlighting the potential of miR-502-3p suppression as a means to enhance synaptic growth and plasticity. These observations align with studies showing that reducing the levels of specific miRNAs can promote dendritic growth and synaptic plasticity (5, 29, 30, 31, 32). This suggests that miR-502-3p plays a critical role in regulating neuronal morphology, and its suppression may be beneficial for synaptic development. The TEM analysis revealed a clear impact of miR-502-3p modulation on synaptic ultrastructure. In the miR-502-3p OE LV group, we observed a significant reduction in the number of mitochondria, shorter mitochondrial length, and fewer presynaptic terminals, suggesting a disruption of synaptic function and energy homeostasis. On the other hand, the miR-502-3p sponge LV group exhibited an increase in synapse number, indicating improved synaptic function and integrity following miR-502-3p suppression. These ultrastructural changes provide further evidence that miR-502-3p plays a crucial role in maintaining synaptic health and function. This finding is particularly important because synaptic number and structure are tightly linked to cognitive function and plasticity (33, 34, 35). The increase in synapse count following miR-502-3p suppression supports the idea that miR-502-3p modulates synaptic structure and function in a manner that could be beneficial for restoring synaptic integrity in neurodegenerative conditions. These results are consistent with research showing that miRNA suppression can reverse synaptic deficits and improve neuronal connectivity (26, 36, 37). Collectively, our results suggest that miR-502-3p plays a critical role in regulating synaptic integrity and plasticity. Overexpression of miR-502-3p impairs synaptic protein expression, dendritic morphology, and synaptic ultrastructure, while suppression of miR-502-3p enhances these features, pointing to its potential as a modulator of synaptic function. These findings contribute to our understanding of the molecular mechanisms underlying synaptic plasticity and offer potential therapeutic avenues for conditions involving synaptic dysfunction. Future studies will be needed to further elucidate the precise molecular targets of miR-502-3p and its broader implications in neurodevelopmental and neurodegenerative disorders.

## CONCLUSION

In conclusion, this study highlights the critical role of miR-502-3p in regulating synaptic structure, function, and plasticity. Through modulation of miR-502-3p expression *via* lentiviral vectors, the study demonstrates significant effects on synaptic protein expression, dendritic morphology, and synaptic ultrastructure, suggesting that miR-502-3p may be a key regulator of synaptic integrity. However, the findings are limited by the use of specific brain region, a single time point of analysis, and potential off-target effects associated with lentiviral-mediated gene manipulation. Future studies should address these limitations by employing more dynamic and comprehensive approaches, including inducible models, multi-regional analyses, and advanced imaging techniques, to further elucidate the molecular mechanisms of miR-502-3p and explore its therapeutic potential in conditions involving synaptic dysfunction.

## ACKNOWLEDGMENTS

The authors are exceedingly grateful to Prof. Rajkumar Lakshmanaswamy, Chair of the Department of Molecular and Translational Medicine, TTUHSC El Paso for the immense research support. We would like to thank our lab members, Ms. Aditi Kulkarni, Mr. Morgan Ogwo, Mr. Davin Devara, Ms. Yogyana and Ms. Angelica.

## AUTHORS CONTRIBUTION

Conceptualization and supervision: SK; experimental performance: BS, DR, GG, RA; LM, KD and SE analysis, interpretation, and validation of data: SK, BS and MAM; writing and original draft preparation: SK and BS; review, editing, and finalization of manuscript: BS, DR, MAM, and SK. All authors have read and agreed to the published version of the manuscript.

## DATA AVAILABILITY

Data will be made available on request.

## REFERENCES

1. Pelucchi S, Gardoni F, Di Luca M, Marcello E. Synaptic dysfunction in early phases of Alzheimer’s Disease. Handb Clin Neurol. 2022;184:417–38.

2. Padmanabhan P, Kneynsberg A, Götz J. Super-resolution microscopy: a closer look at synaptic dysfunction in Alzheimer disease. Nature Reviews Neuroscience. 2021;22(12):723–40.

3. Rivera J, Sharma B, Torres MM, Kumar S. Factors affecting the GABAergic synapse function in Alzheimer’s disease: Focus on microRNAs. Ageing Research Reviews. 2023;92:102123.

4. Li Y-B, Fu Q, Guo M, Du Y, Chen Y, Cheng Y. MicroRNAs: pioneering regulators in Alzheimer’s disease pathogenesis, diagnosis, and therapy. Translational Psychiatry. 2024;14(1):367.

5. Kumar S, Reddy PH. The role of synaptic microRNAs in Alzheimer’s disease. Biochimica et Biophysica Acta (BBA) - Molecular Basis of Disease. 2020;1866(12):165937.

6. Kumar S, Orlov E, Gowda P, Bose C, Swerdlow RH, Lahiri DK, et al. Synaptosome microRNAs regulate synapse functions in Alzheimer’s disease. npj Genomic Medicine. 2022;7(1):47.

7. Wang IF, Ho PC, Tsai KJ. MicroRNAs in Learning and Memory and Their Impact on Alzheimer’s Disease. Biomedicines. 2022;10(8).

8. Song Y, Hu M, Zhang J, Teng Z-q, Chen C. A novel mechanism of synaptic and cognitive impairments mediated via microRNA-30b in Alzheimer’s disease. eBioMedicine. 2019;39:409–21.

9. Zhang Y, Chen H, Li R, Sterling K, Song W. Amyloid β-based therapy for Alzheimer’s disease: challenges, successes and future. Signal Transduction and Targeted Therapy. 2023;8(1):248.

10. Hampel H, Hardy J, Blennow K, Chen C, Perry G, Kim SH, et al. The Amyloid-β Pathway in Alzheimer’s Disease. Molecular Psychiatry. 2021;26(10):5481–503.

11. Taddei RN, Perbet R, Mate de Gerando A, Wiedmer AE, Sanchez-Mico M, Connors Stewart T, et al. Tau Oligomer–Containing Synapse Elimination by Microglia and Astrocytes in Alzheimer Disease. JAMA Neurology. 2023;80(11):1209–21.

12. Colom-Cadena M, Davies C, Sirisi S, Lee J-E, Simzer EM, Tzioras M, et al. Synaptic oligomeric tau in Alzheimer&#x2019;s disease &#x2014; A potential culprit in the spread of tau pathology through the brain. Neuron. 2023;111(14):2170–83.e6.

13. Devara D, Sharma B, Goyal G, Rodarte D, Kulkarni A, Tinu N, et al. MiRNA-501-3p and MiRNA-502-3p: A Promising Biomarker Panel for Alzheimer’s Disease. bioRxiv. 2025:2025.01.09.632227.

14. Sharma B, Torres MM, Rodriguez S, Gangwani L, Kumar S. MicroRNA-502-3p regulates GABAergic synapse function in hippocampal neurons. Neural Regen Res. 2024;19(12):2698–707.

15. Limon A, Reyes-Ruiz JM, Miledi R. Loss of functional GABA_A_ receptors in the Alzheimer diseased brain. Proceedings of the National Academy of Sciences. 2012;109(25):10071–6.

16. Murari G, Liang DR-S, Ali A, Chan F, Mulder-Heijstra M, Verhoeff NPLG, et al. Prefrontal GABA Levels Correlate with Memory in Older Adults at High Risk for Alzheimer’s Disease. Cerebral Cortex Communications. 2020;1(1).

17. Cutler AJ, Mattingly GW, Maletic V. Understanding the mechanism of action and clinical effects of neuroactive steroids and GABAergic compounds in major depressive disorder. Translational Psychiatry. 2023;13(1):228.

18. Goetz T, Arslan A, Wisden W, Wulff P. GABA(A) receptors: structure and function in the basal ganglia. Prog Brain Res. 2007;160:21–41.

19. Yan C, Mercaldo V, Jacob AD, Kramer E, Mocle A, Ramsaran AI, et al. Higher-order interactions between hippocampal CA1 neurons are disrupted in amnestic mice. Nature Neuroscience. 2024;27(9):1794–804.

20. Jeong N, Singer AC. Learning from inhibition: Functional roles of hippocampal CA1 inhibition in spatial learning and memory. Current Opinion in Neurobiology. 2022;76:102604.

21. Masurkar AV. Towards a circuit-level understanding of hippocampal CA1 dysfunction in Alzheimer’s disease across anatomical axes. J Alzheimers Dis Parkinsonism. 2018;8(1).

22. Devara D, Choudhary Y, Kumar S. Role of MicroRNA-502-3p in Human Diseases. Pharmaceuticals (Basel). 2023;16(4).

23. Paxinos AO, Franklin Ma KBJ. Paxinos and Franklin’s the Mouse Brain in Stereotaxic Coordinates. Fifth edition. ed. Chantilly: Elsevier Science & Technology; 2019.

24. Pan J, Qu M, Li Y, Wang L, Zhang L, Wang Y, et al. MicroRNA-126-3p/-5p Overexpression Attenuates Blood-Brain Barrier Disruption in a Mouse Model of Middle Cerebral Artery Occlusion. Stroke. 2020;51(2):619–27.

25. Du M, Wu C, Yu R, Cheng Y, Tang Z, Wu B, et al. A novel circular RNA, circIgfbp2, links neural plasticity and anxiety through targeting mitochondrial dysfunction and oxidative stress-induced synapse dysfunction after traumatic brain injury. Molecular Psychiatry. 2022;27(11):4575–89.

26. Barros-Viegas AT, Carmona V, Ferreiro E, Guedes J, Cardoso AM, Cunha P, et al. miRNA- 31 Improves Cognition and Abolishes Amyloid-β Pathology by Targeting APP and BACE1 in an Animal Model of Alzheimer’s Disease. Molecular Therapy - Nucleic Acids. 2020;19:1219–36.

27. Sun K, Guo C, Deng H-j, Dong J-q, Lei S-t, Li G-x. Construction of lentivirus-based inhibitor of hsa-microRNA-338-3p with specific secondary structure. Acta Pharmacologica Sinica. 2013;34(1):167–75.

28. Kapplingattu SV, Bhattacharya S, Adlakha YK. MiRNAs as major players in brain health and disease: current knowledge and future perspectives. Cell Death Discovery. 2025;11(1):7.

29. Martins HC, Schratt G. MicroRNA-dependent control of neuroplasticity in affective disorders. Translational Psychiatry. 2021;11(1):263.

30. Yoshino Y, Roy B, Dwivedi Y. Differential and unique patterns of synaptic miRNA expression in dorsolateral prefrontal cortex of depressed subjects. Neuropsychopharmacology. 2021;46(5):900–10.

31. Parkins EV, Brager DH, Rymer JK, Burwinkel JM, Rojas D, Tiwari D, et al. Mir324 knockout regulates the structure of dendritic spines and impairs hippocampal long-term potentiation. Scientific Reports. 2023;13(1):21919.

32. Cohen JE, Lee PR, Chen S, Li W, Fields RD. MicroRNA regulation of homeostatic synaptic plasticity. Proceedings of the National Academy of Sciences. 2011;108(28):11650–5.

33. Santuy A, Tomás-Roca L, Rodríguez J-R, González-Soriano J, Zhu F, Qiu Z, et al. Estimation of the number of synapses in the hippocampus and brain-wide by volume electron microscopy and genetic labeling. Scientific Reports. 2020;10(1):14014.

34. Colom-Cadena M, Spires-Jones T, Zetterberg H, Blennow K, Caggiano A, DeKosky ST, et al. The clinical promise of biomarkers of synapse damage or loss in Alzheimer’s disease. Alzheimer’s Research & Therapy. 2020;12(1):21.

35. Taddei RN, E. Duff K. Synapse vulnerability and resilience underlying Alzheimer&#x2019;s disease. eBioMedicine. 2025;112.

36. Rodrigues B, Leitão RA, Santos M, Trofimov A, Silva M, Inácio ÂS, et al. MiR-186-5p inhibition restores synaptic transmission and neuronal network activity in a model of chronic stress. Molecular Psychiatry. 2025;30(3):1034–46.

37. Li Y, Fan C, Wang L, Lan T, Gao R, Wang W, et al. MicroRNA-26a-3p rescues depression-like behaviors in male rats via preventing hippocampal neuronal anomalies. The Journal of Clinical Investigation. 2021;131(16).

