## Supplementary information file 1 for "Modulation of microRNA-502-3p significantly influences synaptic activity, dendritic spine density and mitochondrial morphology in the mice brain"

### Vector Summary

|  |  |
| --- | --- |
| Vector ID | VB010000-0009mxc |
| Vector Name | pLV[shRNA]-EGFP/Puro-U6>Scramble_shRNA |
| Vector Size | 8347 bp |
| Viral Genome Size | 4872 bp |
| Vector Type | Mammalian shRNA Knockdown Lentiviral Vector |
| Inserted shRNA | Scramble_shRNA#1 |
| Target Sequence | CCTAAGGTTAAGTCGCCCTCG |
| Inserted Marker | EGFP:T2A:Puro |
| Plasmid Copy Number | High |
| Antibiotic Resistance | Ampicillin |
| Cloning Host | VB UltraStable (or alternative strain) |

### Vector Map

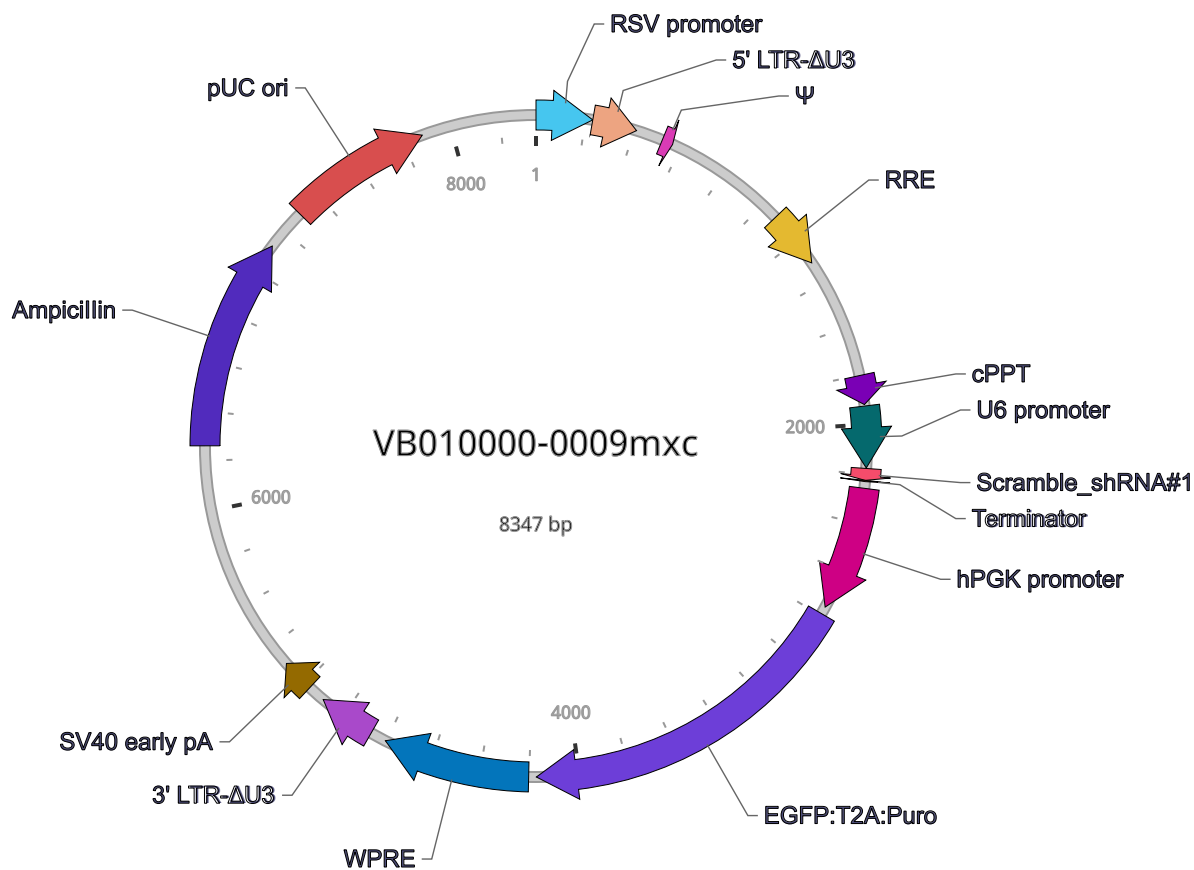

### Vector Components

| Name | Position | Size (bp) | Type | Description | Application notes |
| --- | --- | --- | --- | --- | --- |
| RSV promoter | 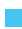 1-229 | 229 | Promoter | Rous sarcoma virus enhancer/promoter | Strong promoter; drives transcription of viral RNA in packaging cells. |
| 5' LTR-ΔU3 | 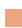 230-410 | 181 | LTR | Truncated HIV-1 5' long terminal repeat | Allows transcription of viral RNA and its packaging into virus. |
| Ψ | 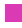 521-565 | 45 | Miscellaneous | HIV-1 packaging signal | Allows packaging of viral RNA into virus. |
| RRE | 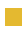 1075-1308 | 234 | Miscellaneous | HIV-1 Rev response element | Rev protein binding site that allows Rev-dependent nuclear export of viral RNA during viral packaging. |
| cPPT | 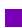 1803-1920 | 118 | Miscellaneous | Central polypurine tract | Facilitates the nuclear import of HIV-1 cDNA through a central DNA flap. |
| U6 promoter | 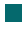 1927-2175 | 249 | Promoter | Human U6 small nuclear 1 promoter | Pol III promoter; drives expression of small RNAs. |
| **Scramble_shRNA#1** | 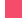 2178-2225 | 48 | shRNA | *None* | Target: None in human and mouse. |
| Terminator | 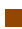 2226-2230 | 5 | terminator | Pol III transcription terminator | Allows transcription termination of small RNA transcribed by Pol III RNA polymerase. |
| hPGK promoter | 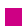 2258-2762 | 505 | Promoter | Human phosphoglycerate kinase 1 promoter | Medium-strength promoter. |
| **EGFP:T2A:Puro** | 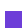 2793-4172 | 1380 | CDS | EGFP and Puro linked by T2A | Allows cells to be visualized by green fluorescence and resistant to puromycin. |
| WPRE | 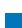 4204-4801 | 598 | Miscellaneous | Woodchuck hepatitis virus posttranscriptional regulatory element | Enhances virus stability in packaging cells, leading to higher titer of packaged virus; enhances higher expression of transgenes. |
| 3' LTR-ΔU3 | ■ 4867-5101 | 235 | LTR | Truncated HIV-1 3' long terminal repeat | Allows packaging of viral RNA into virus; self-inactivates the 5' LTR by a copying mechanism during viral genome integration; contains polyadenylation signal for transcription termination. |
| SV40 early pA | ■ 5174-5308 | 135 | PolyA_signal | Simian virus 40 early polyadenylation signal | Allows transcription termination and polyadenylation of mRNA transcribed by Pol II RNA polymerase. |
| Ampicillin | ■ 6262-7122 | 861 | CDS | Ampicillin resistance gene | Allows E. coli to be resistant to ampicillin. |
| pUC ori | ■ 7293-7881 | 589 | Rep_origin | pUC origin of replication | Facilitates plasmid replication in E. coli; regulates high-copy plasmid number (500-700). |

Note: Components added by user are listed in **bold red** text.

### Vector Sequence

```
1  AATGTAGTCT TATGCAATAC TCTTGTAGTC TTGCAACATG GTAACGATGA GTTAGCAACA TGCCTTACAA GGAGAGAAAA
81 AGCACCGTGC ATGCCGATTG GTGGAAGTAA GGTGGTACGA TCGTGCCTTA TTAGGAAGGC AACAGACGGG TCTGACATGG
161 ATTGGACGAA CCACTGAATT GCCGCATTGC AGAGATATTG TATTTAAGTG CCTAGCTCGA TACATAAACG GGTCTCTCTG
241 GTTAGACCAG ATCTGAGCCT GGGAGCTCTC TGGCTAACTA GGGAACCCAC TGCTTAAGCC TCAATAAAGC TTGCCTTGAG
321 TGCTTCAAGT AGTGTGTGCC CGTCTGTTGT GTGACTCTGG TAACTAGAGA TCCCTCAGAC CCTTTTAGTC AGTGTGGAAA
401 ATCTCTAGCA GTGGCGCCCG AACAGGGACT TGAAAGCGAA AGGGAAACCA GAGGAGCTCT CTCGACGCAG GACTCGGCTT
481 GCTGAAGCGC GCACGGCAAG AGGCGAGGGG CGGCGACTGG TGAGTACGCC AAAAATTTTG ACTAGCGGAG GCTAGAAGGA
561 GAGAGATGGG TGCGAGAGCG TCAGTATTAA GCGGGGGAGA ATTAGATCGC GATGGGAAAA AATTCGGTTA AGGCCAGGGG
641 GAAAGAAAAA ATATAAATTA AAACATATAG TATGGGCAAG CAGGGAGCTA GAACGATTCG CAGTTAATCC TGGCCTGTTA
721 GAAACATCAG AAGGCTGTAG ACAAATACTG GGACAGCTAC AACCATCCCT TCAGACAGGA TCAGAAGAAC TTAGATCATT
801 ATATAATACA GTAGCAACCC TCTATTGTGT GCATCAAAGG ATAGAGATAA AAGACACCAA GGAAGCTTTA GACAAGATAG
881 AGGAAGAGCA AAACAAAAGT AAGACCACCG CACAGCAAGC GGCCGCTGAT CTTCAGACCT GGAGGAGGAG ATATGAGGGA
961 CAATTGGAGA AGTGAATTAT ATAAATATAA AGTAGTAAAA ATTGAACCAT TAGGAGTAGC ACCCACCAAG GCAAAGAGAA
1041 GAGTGGTGCA GAGAGAAAAA AGAGCAGTGG GAATAGGAGC TTTGTTCCTT GGGTTCTTGG GAGCAGCAGG AAGCACTATG
1121 GGCGCAGCGT CAATGACGCT GACGGTACAG GCCAGACAAT TATTGTCTGG TATAGTGCAG CAGCAGAACA ATTTGCTGAG
1201 GGCTATTGAG GCGCAACAGC ATCTGTTGCA ACTCACAGTC TGGGGCATCA AGCAGCTCCA GGCAAGAATC CTGGCTGTGG
1281 AAAGATACCT AAAGGATCAA CAGCTCCTGG GGATTTGGGG TTGCTCTGGA AAACTCATTT GCACCACTGC TGTGCCTTGG
1361 AATGCTAGTT GGAGTAATAA ATCTCTGGAA CAGATTTGGA ATCACACGAC CTGGATGGAG TGGGACAGAG AAATTAACAA
```

```
1441 TTACACAAGC TTAATACACT CCTTAATTGA AGAATCGCAA AACCAGCAAG AAAAGAATGA ACAAGAATTA TTGGAATTAG
1521 ATAAATGGGC AAGTTTGTGG AATTGGTTTA ACATAACAAA TTGGCTGTGG TATATAAAAT TATTCATAAT GATAGTAGGA
1601 GGCTTGGTAG GTTTAAGAAT AGTTTTTGCT GTACTTTCTA TAGTGAATAG AGTTAGGCAG GGATATTCAC CATTATCGTT
1681 TCAGACCCAC CTCCCAACCC CGAGGGGACC CGACAGGCCC GAAGGAATAG AAGAAGAAGG TGGAGAGAGA GACAGAGACA
1761 GATCCATTCG ATTAGTGAAC GGATCTCGAC GGTATCGCTA GCTTTTAAAA GAAAAGGGGG GATTGGGGGG TACAGTGCAG
1841 GGGAAAGAAT AGTAGACATA ATAGCAACAG ACATACAAAC TAAAGAATTA CAAAAACAAA TTACAAAAAT TCAAAATTTT
1921 ACTAGTGAGG GCCTATTTCC CATGATTCCT TCATATTTGC ATATACGATA CAAGGCTGTT AGAGAGATAA TTGGAATTAA
2001 TTTGACTGTA AACACAAAGA TATTAGTACA AAATACGTGA CGTAGAAAGT AATAATTTCT TGGGTAGTTT GCAGTTTTAA
2081 AATTATGTTT TAAAATGGAC TATCATATGC TTACCGTAAC TTGAAAGTAT TTCGATTTCT TGGCTTTATA TATCTTGTGG
2161 AAAGGACGAA ACACCGGCCT AAGGTTAAGT CGCCCTCGCT CGAGCGAGGG CGACTTAACC TTAGGTTTTT GAATTCCAAC
2241 TTTGTATAGA AAAGTTGGGG TTGCGCCTTT TCCAAGGCAG CCCTGGGTTT GCGCAGGGAC GCGGCTGCTC TGGGCGTGGT
2321 TCCGGGAAAC GCAGCGGCGC CGACCCTGGG TCTCGCACAT TCTTCACGTC CGTTCGCAGC GTCACCCGGA TCTTCGCCGC
2401 TACCCTTGTG GGCCCCCCGG CGACGCTTCC TGCTCCGCCC CTAAGTCGGG AAGGTTCCTT GCGGTTCGCG GCGTGCCGGA
2481 CGTGACAAAC GGAAGCCGCA CGTCTCACTA GTACCCTCGC AGACGGACAG CGCCAGGGAG CAATGGCAGC GCGCCGACCG
2561 CGATGGGCTG TGGCCAATAG CGGCTGCTCA GCAGGGCGCG CCGAGAGCAG CGGCCGGGAA GGGGCGGTGC GGGAGGCGGG
2641 GTGTGGGGCG GTAGTGTGGG CCCTGTTCCT GCCCGCGCGG TGTTCCGCAT TCTGCAAGCC TCCGGAGCGC ACGTCGGCAG
2721 TCGGCTCCCT CGTTGACCGA ATCACCGACC TCTCTCCCCA GGCAAGTTTG TACAAAAAAG CAGGCTGCCA CCATGGTGAG
2801 CAAGGGCGAG GAGCTGTTCA CCGGGGTGGT GCCCATCCTG GTCGAGCTGG ACGGCGACGT AAACGGCCAC AAGTTCAGCG
2881 TGTCCGGCGA GGGCGAGGGC GATGCCACCT ACGGCAAGCT GACCCTGAAG TTCATCTGCA CCACCGGCAA GCTGCCCGTG
2961 CCCTGGCCCA CCCTCGTGAC CACCCTGACC TACGGCGTGC AGTGCTTCAG CCGCTACCCC GACCACATGA AGCAGCACGA
3041 CTTCTTCAAG TCCGCCATGC CCGAAGGCTA CGTCCAGGAG CGCACCATCT TCTTCAAGGA CGACGGCAAC TACAAGACCC
3121 GCGCCGAGGT GAAGTTCGAG GGCGACACCC TGGTGAACCG CATCGAGCTG AAGGGCATCG ACTTCAAGGA GGACGGCAAC
3201 ATCCTGGGGC ACAAGCTGGA GTACAACTAC AACAGCCACA ACGTCTATAT CATGGCCGAC AAGCAGAAGA ACGGCATCAA
3281 GGTGAACTTC AAGATCCGCC ACAACATCGA GGACGGCAGC GTGCAGCTCG CCGACCACTA CCAGCAGAAC ACCCCCATCG
3361 GCGACGGCCC CGTGCTGCTG CCCGACAACC ACTACCTGAG CACCCAGTCC GCCCTGAGCA AAGACCCCAA CGAGAAGCGC
3441 GATCACATGG TCCTGCTGGA GTTCGTGACC GCCGCCGGGA TCACTCTCGG CATGGACGAG CTGTACAAGG GCTCCGGAGA
3521 GGGCAGGGGA AGTCTTCTAA CATGCGGGGA CGTGGAGGAA AATCCCGGCC CCATGACCGA GTACAAGCCC ACGGTGCGCC
3601 TCGCCACCCG CGACGACGTC CCCAGGGCCG TACGCACCCT CGCCGCCGCG TTCGCCGACT ACCCCGCCAC GCGCCACACC
3681 GTCGATCCGG ACCGCCACAT CGAGCGGGTC ACCGAGCTGC AAGAACTCTT CCTCACGCGC GTCGGGCTCG ACATCGGCAA
3761 GGTGTGGGTC GCGGACGACG GCGCCGCGGT GGCGGTCTGG ACCACGCCGG AGAGCGTCGA AGCGGGGGCG GTGTTCGCCG
3841 AGATCGGCCC GCGCATGGCC GAGTTGAGCG GTTCCCGGCT GGCCGCGCAG CAACAGATGG AAGGCCTCCT GGCGCCGCAC
3921 CGGCCCAAGG AGCCCGCGTG GTTCCTGGCC ACCGTCGGCG TCTCGCCCGA CCACCAGGGC AAGGGTCTGG GCAGCGCCGT
4001 CGTGCTCCCC GGAGTGGAGG CGGCCGAGCG CGCCGGGGTG CCCGCCTTCC TGGAGACCTC CGCGCCCCGC AACCTCCCCT
4081 TCTACGAGCG GCTCGGCTTC ACCGTCACCG CCGACGTCGA GGTGCCCGAA GGACCGCGCA CCTGGTGCAT GACCCGCAAG
4161 CCCGGTGCCT GAACCCAGCT TTCTTGTACA AAGTGGTGGT ACCCGATAAT CAACCTCTGG ATTACAAAAT TTGTGAAAGA
4241 TTGACTGGTA TTCTTAACTA TGTTGCTCCT TTTACGCTAT GTGGATACGC TGCTTTAATG CCTTTGTATC ATGCTATTGC
4321 TTCCCGTATG GCTTTCATTT TCTCCTCCTT GTATAAATCC TGGTTGCTGT CTCTTTATGA GGAGTTGTGG CCCGTTGTCA
4401 GGCAACGTGG CGTGGTGTGC ACTGTGTTTG CTGACGCAAC CCCCACTGGT TGGGGCATTG CCACCACCTG TCAGCTCCTT
4481 TCCGGGACTT TCGCTTTCCC CCTCCCTATT GCCACGGCGG AACTCATCGC CGCCTGCCTT GCCCGCTGCT GGACAGGGGC
4561 TCGGCTGTTG GGCACTGACA ATTCCGTGGT GTTGTCGGGG AAGCTGACGT CCTTTCCATG GCTGCTCGCC TGTGTTGCCA
4641 CCTGGATTCT GCGCGGGACG TCCTTCTGCT ACGTCCCTTC GGCCCTCAAT CCAGCGGACC TTCCTTCCCG CGGCCTGCTG
4721 CCGGCTCTGC GGCCTCTTCC GCGTCTTCGC CTTCGCCCTC AGACGAGTCG GATCTCCCTT TGGGCCGCCT CCCCGCATCG
4801 GCTTTAAGAC CAATGACTTA CAAGGCAGCT GTAGATCTTA GCCACTTTTT AAAAGAAAAG GGGGGACTGG AAGGGCTAAT
4881 TCACTCCCAA CGAAGACAAG ATCTGCTTTT TGCTTGTACT GGGTCTCTCT GGTTAGACCA GATCTGAGCC TGGGAGCTCT
4961 CTGGCTAACT AGGGAACCCA CTGCTTAAGC CTCAATAAAG CTTGCCTTGA GTGCTTCAAG TAGTGTGTGC CCGTCTGTTG
5041 TGTGACTCTG GTAAC TAGAG ATCCCTCAGA CCCTTTTAGT CAGTGTGGAA AATCTCTAGC AGTAGTAGTT CATGTCATCT
5121 TATTATTCAG TATTTATAAC TTGCAAAGAA ATGAATATCA GAGAGTGAGA GGAACTTGTT TATTGCAGCT TATAATGGTT
5201
```

```
5281  ACAAATAAAG  CAATAGCATC  ACAAATTTCA  CAAATAAAGC  ATTTTTTTCA  CTGCATTCTA  GTTGTGGTTT  GTCCAAACTC
5361  ATCAATGTAT  CTTATCATGT  CTGGCTCTAG  CTATCCCGCC  CCTAACTCCG  CCCATCCCGC  CCCTAACTCC  GCCCAGTTCC
5441  GCCCATTCTC  CGCCCCATGG  CTGACTAATT  TTTTTTATTT  ATGCAGAGGC  CGAGGCCGCC  TCGGCCTCTG  AGCTATTCCA
5521  GAAGTAGTGA  GGAGGCTTTT  TTGGAGGCCT  AGGGACGTAC  CCAATTCGCC  CTATAGTGAG  TCGTATTACG  CGCGCTCACT
5601  GGCCGTCGTT  TTACAACGTC  GTGACTGGGA  AAACCCTGGC  GTTACCCAAC  TTAATCGCCT  TGCAGCACAT  CCCCTTTTCG
5681  CCAGCTGGCG  TAATAGCGAA  GAGGCCCGCA  CCGATCGCCC  TTCCAACAG  TTGCGCAGCC  TGAATGGCGA  ATGGGACGCG
5761  CCCTGTAGCG  GCGCATTAAG  CGCGGCGGGT  GTGGTGGTTA  CGCGCAGCGT  GACCGCTACA  CTTGCCAGCG  CCCTAGCGCC
5841  CGCTCCTTTC  GCTTCTTCC  CTTCTTTCT  CGCCACGTTT  GCCGGCTTTC  CCCGTCAAGC  TCTAAATCGG  GGGCTCCCTT
5921  TAGGGTCCG  ATTTAGTGCT  TTACGGCACC  TCGACCCCAA  AAAACTTGAT  TAGGGTGATG  GTTCACGTAG  TGGGCCATCG
6001  CCCTGATAGA  CGGTTTTTCG  CCCTTTGACG  TTGGAGTCCA  CGTTCCTTAA  TAGTGGAATC  TTGTTCCTAA  CTGGAACAAC
6081  ACTCAACCT  ATCTCGGTCT  ATTCTTTTGA  TTTATAAGGG  ATTTTGCCGA  TTTTCGCCTA  TTGGTTAAAA  AATGAGCTGA
6161  TTTAACAAAA  ATTTAACGCG  AATTTTAAAC  AAATATTAAC  GCTTACAATT  TAGGTGGCAC  TTTTCGGGGA  AATGTGCGCG
6241  GAACCCCTAT  TTGTTTATTT  TTCTAAATAC  ATTCAAATAT  GTATCCGCTC  ATGAGACAA  AACCCTGATA  AATGCTTCAA
6321  TAATATTGAA  AAAGGAAGAG  TATGAGTATT  CAACATTTCC  GTGTCGCCCT  TATTCCTTTT  TTTGCGGCAT  TTTGCCTTCC
6401  TGTTTTTGCT  CACCCAGAAA  CGCTGGTGAA  AGTAAAAGAT  GCTGAAGATC  AGTTGGGTGC  ACGAGTGGGT  TACATCGAAC
6481  TGGATCTCAA  CAGCGGTAAG  ATCCTTGAGA  GTTTTCGCCC  CGAAGAACGT  TTTCCAATGA  TGAGCACTTT  TAAAGTTCTG
6561  CTATGTGGCG  CGGTATTATC  CCGTATTGAC  GCCGGGCAAG  AGCAACTCGG  TCGCCGCATA  CACTATTCTC  AGAATGACTT
6641  GGTGAGTAC  TCACCAGTCA  CAGAAAAGCA  TCTTACGGAT  GGCATGACAG  TAAGAGAATT  ATGCAGTGCT  GCCATAACCA
6721  TGAGTGATAA  CACTGCGGCC  AACTTACTTC  TGACAACGAT  CGGAGGACCG  AAGGAGCTAA  CCGCTTTTTT  GCACAACATG
6801  GGGGATCATG  TAACTCGCCT  TGATCGTTGG  GAACCGGAGC  TGAATGAAGC  CATACCAAAC  GACGAGCGTG  ACACCACGAT
6881  GCCTGTAGCA  ATGGCAACAA  CGTTGCACAA  ACTATTAACT  GGCGAACTAC  TTACTCTAGC  TTCCCGGCAA  CAATTAATAG
6961  ACTGGATGGA  GGGCGATAAA  GTTGCAGGAC  CACTTCTGCG  CTCGGCCCTT  CCGGCTGGCT  GGTATTATGC  TGATAAATCT
7041  GGAGCCGGTG  AGCGTGGGTC  TCGCGGTATC  ATTGCAGCAC  TGGGGCCAGA  TGTAAGCCG  TCCCGTATCG  TAGTTATCTA
7121  CACGACGGGG  AGTCAGGCAA  CTATGGATGA  ACGAAATAGA  CAGATCGCTG  AGATAGGTGC  CTCACTGATT  AAGCATTGGT
7201  AACTGTCAGA  CCAAGTTTAC  TCATATATAC  TTTAGATTGA  TTTAAAACCT  CATTTTAAAT  TTTAAAGGAT  CTAGGTGAAG
7281  ATCCTTTTTG  ATAATCTCAT  GACCAAAATC  CCTTAACGTG  AGTTTTCGTT  CCACTGAGCG  TCAGACCCCG  TAGAAAAGAT
7361  CAAAGGATCT  TCTTGAGATC  CTTTTTTTCT  GCGCGTAATC  TGCTGCTTGC  AAACAAAAAA  ACCACCGCTA  CCAGCGGTGG
7441  TTTGTTTGCC  GGATCAAGAG  CTACCAACTC  TTTTCCGAA  GGTAAGTGGC  TTCAGCAGAG  CGCAGATACC  AAATACTGTT
7521  CTTCTAGTGT  AGCCGTAGTT  AGGCCACCAC  TTCAAGAACT  CTGTAGCACC  GCCTACATAC  CTCGCTCTGC  TAATCCTGTT
7601  ACCAGTGGCT  GCTGCCAGTG  GCGATAAGTC  GTGTCTTACC  GGGTTGGACT  CAAGACGATA  GTTACCGGAT  AAGGCGCAGC
7681  GGTTCGGGCT  AACGGGGGGT  TCGTGCACAC  AGCCCAGCTT  GGAGCGAACG  ACCTACACCG  AACTGAGATA  CCTACAGCGT
7761  GAGCTATGAG  AAAGCGCCAC  GCTTCCCGAA  GAGAGAAAGG  CGGACAGGTA  TCCGGTAAGC  GGCAGGGTCG  GAACAGGAGA
7841  GCGCACGAGG  GAGCTTCCAG  GGGGAAACGC  CTGGTATCTT  TATAGTCCTG  TCGGGTTTCG  CCACCTCTGA  CTTGAGCGTC
7921  GATTTTGTG  ATGCTCGTCA  GGGGGGCGGA  GCCTATGGAA  AAACGCCAGC  AACGCGGCCT  TTTTACGGTT  CCTGGCCTTT
8001  TGCTGGCCTT  TTGCTCACAT  GTTCTTTCCT  GCGTTATCCC  CTGATTCTGT  GGATAACCGT  ATTACCGCCT  TTGAGTGAGC
8081  TGATACCGCT  CGCCGCAGCC  GAACGACCGA  GCGCAGCGAG  TCAGTGAGCG  AGGAAGCGGA  AGAGCGCCCA  ATACGCAAAC
8161  CGCCTCTCCC  CGCGCGTTGG  CCGATTCAAT  AATGCAGCTG  GCACGACAGG  TTTCCCGACT  GGAAGCGGGG  CAGTGAGCGC
8241  AACGCAATTA  ATGTGAGTTA  GCTCACTCAT  TAGGCACCCC  AGGCTTTACA  CTTTATGCTT  CCGGCTCGTA  TGTTGTGTGG
8321  AATTGTGAGC  GGATAACAAT  TTCACACAGG  AAACAGCTAT  GACCATGATT  ACGCCAAGCG  CGCAATTAAC  CCTCACTAAA
      GGGAACAAAA  GCTGGAGCTG  CAAGCTT
```

### Validation by Restriction Enzyme Digestion

| Restriction Enzymes | Cutting Sites | DNA Fragments (bp) |
| --- | --- | --- |
| XhoI | 2200 | 8347 |
| ApaLI | 4418, 6378, 7624 | 1960, 1246, 5141 |
| ApaLI+XhoI | 2200, 4418, 6378, 7624 | 2218, 1960, 1246, 2923 |

### Vector Summary

|  |  |
| --- | --- |
| Vector ID | VB231005-1051gew |
| Vector Name | pLV[shRNA]-EGFP-U6>{hsa-microRNA-502-3p} |
| Vector Size | 7689 bp |
| Viral Genome Size | 4214 bp |
| Vector Type | Mammalian shRNA Knockdown Lentiviral Vector |
| Inserted shRNA | {hsa-microRNA-502-3p} |
| Target Sequence | TGAATCCTTGCCCAGGTGCATT |
| Inserted Marker | EGFP |
| Plasmid Copy Number | High |
| Antibiotic Resistance | Ampicillin |
| Cloning Host | VB UltraStable (or alternative strain) |

### Vector Map

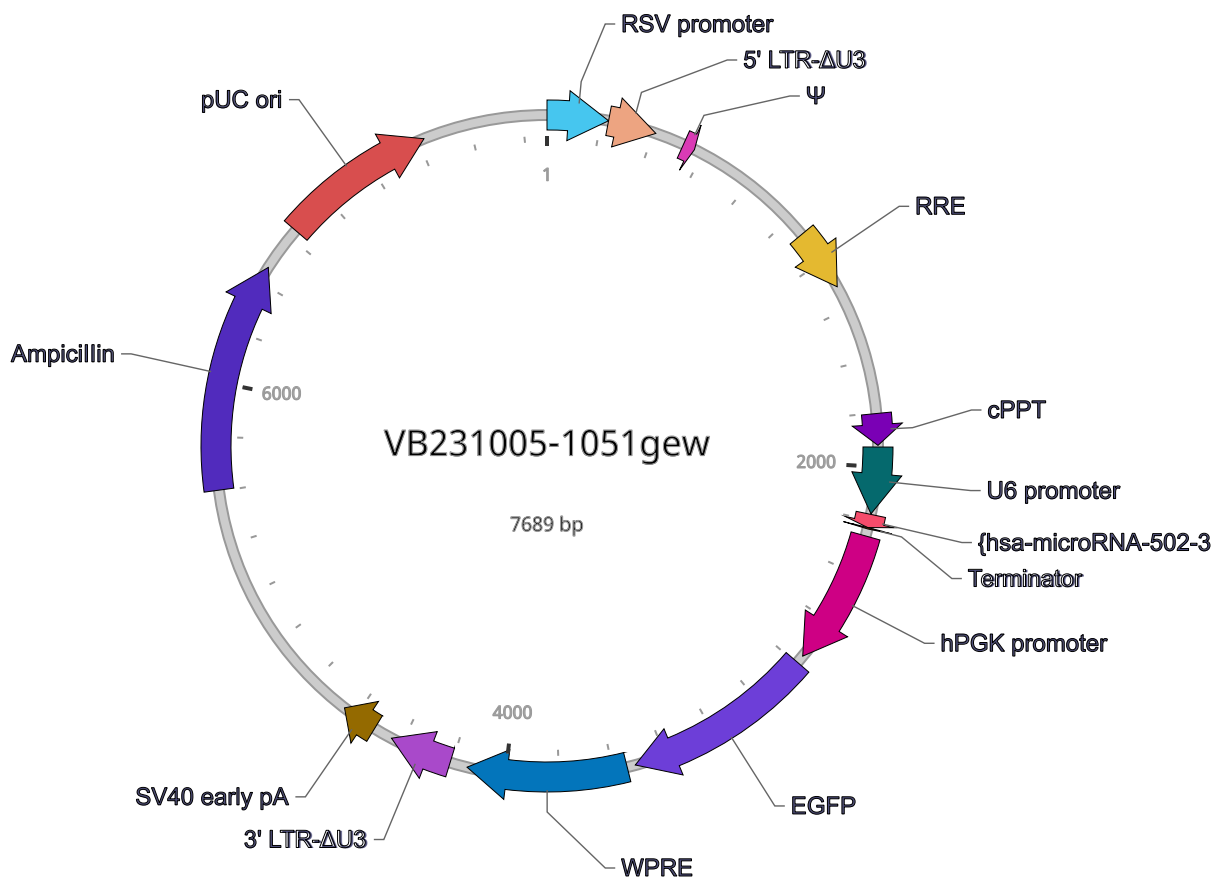

### Vector Components

| Name | Position | Size (bp) | Type | Description | Application notes |
| --- | --- | --- | --- | --- | --- |
| RSV promoter | 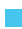 1-229 | 229 | Promoter | Rous sarcoma virus enhancer/promoter | Strong promoter; drives transcription of viral RNA in packaging cells. |
| 5' LTR-ΔU3 | 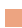 230-410 | 181 | LTR | Truncated HIV-1 5' long terminal repeat | Allows transcription of viral RNA and its packaging into virus. |
| Ψ | 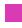 521-565 | 45 | Miscellaneous | HIV-1 packaging signal | Allows packaging of viral RNA into virus. |
| RRE | 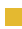 1075-1308 | 234 | Miscellaneous | HIV-1 Rev response element | Rev protein binding site that allows Rev-dependent nuclear export of viral RNA during viral packaging. |
| cPPT | 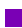 1803-1920 | 118 | Miscellaneous | Central polypurine tract | Facilitates the nuclear import of HIV-1 cDNA through a central DNA flap. |
| U6 promoter | 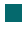 1927-2175 | 249 | Promoter | Human U6 small nuclear 1 promoter | Pol III promoter; drives expression of small RNAs. |
| {**hsa-microRNA-502-3p**} | 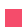 2178-2227 | 50 | shRNA | *None* | *None* |
| Terminator | 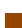 2228-2232 | 5 | terminator | Pol III transcription terminator | Allows transcription termination of small RNA transcribed by Pol III RNA polymerase. |
| hPGK promoter | 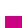 2260-2764 | 505 | Promoter | Human phosphoglycerate kinase 1 promoter | Medium-strength promoter. |
| **EGFP** | 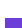 2795-3514 | 720 | CDS | Enhanced green fluorescent protein; codon optimized based on a variant of wild type GFP from the jellyfish *Aequorea victoria* | Commonly used green fluorescent protein; ranked high in brightness, photostability and pH stability among all fluorescent proteins. |
| WPRE | ■ 3546-4143 | 598 | Miscellaneous | Woodchuck hepatitis virus posttranscriptional regulatory element | Enhances virus stability in packaging cells, leading to higher titer of packaged virus; enhances higher expression of transgenes. |
| 3' LTR-ΔU3 | ■ 4209-4443 | 235 | LTR | Truncated HIV-1 3' long terminal repeat | Allows packaging of viral RNA into virus; self-inactivates the 5' LTR by a copying mechanism during viral genome integration; contains polyadenylation signal for transcription termination. |
| SV40 early pA | ■ 4516-4650 | 135 | PolyA_signal | Simian virus 40 early polyadenylation signal | Allows transcription termination and polyadenylation of mRNA transcribed by Pol II RNA polymerase. |
| Ampicillin | ■ 5604-6464 | 861 | CDS | Ampicillin resistance gene | Allows E. coli to be resistant to ampicillin. |
| pUC ori | ■ 6635-7223 | 589 | Rep_origin | pUC origin of replication | Facilitates plasmid replication in E. coli; regulates high-copy plasmid number (500-700). |

**Note:** Components added by user are listed in **red** text.

### Vector Sequence

```
1  AATGTAGTCT TATGCAATAC TCTTGTAGTC TTGCAACATG GTAACGATGA GTTAGCAACA TGCCTTACAA GGAGAGAAAA
81 AGCACCGTGC ATGCCGATTG GTGGAAGTAA GGTGGTACGA TCGTGCCTTA TTAGGAAGGC AACAGACGGG TCTGACATGG
161 ATTGGACGAA CCACTGAATT GCCGCATTGC AGAGATATTG TATTTAAGTG CCTAGCTCGA TACATAAACG GGTCTCTCTG
241 GTTAGACCAG ATCTGAGCCT GGGAGCTCTC TGGCTAACTA GGGAACCCAC TGCTTAAGCC TCAATAAAGC TTGCCTTGAG
321 TGCTTCAAGT AGTGTGTGCC CGTCTGTTGT GTGACTCTGG TAACTAGAGA TCCCTCAGAC CCTTTTAGTC AGTGTGGAAA
401 ATCTCTAGCA GTGGCGCCCG AACAGGGACT TGAAAGCGAA AGGGAAACCA GAGGAGCTCT CTCGACGCAG GACTCGGCTT
481 GCTGAAGCGC GCACGGCAAG AGGCGAGGGG CGGCGACTGG TGAGTACGCC AAAAATTTTG ACTAGCGGAG GCTAGAAGGA
561 GAGAGATGGG TGCGAGAGCG TCAGTATTAA GCGGGGGAGA ATTAGATCGC GATGGGAAAA AATTCGGTTA AGGCCAGGGG
641 GAAAGAAAAA ATATAAATTA AAACATATAG TATGGGCAAG CAGGGAGCTA GAACGATTCG CAGTTAATCC TGGCCTGTTA
721 GAAACATCAG AAGGCTGTAG ACAAATACTG GGACAGCTAC AACCATCCCT TCAGACAGGA TCAGAAGAAC TTAGATCATT
801 ATATAATACA GTAGCAACCC TCTATTGTGT GCATCAAAGG ATAGAGATAA AAGACACCAA GGAAGCTTTA GACAAGATAG
881 AGGAAGAGCA AAACAAAAGT AAGACCACCG CACAGCAAGC GGCCGCTGAT CTTCAGACCT GGAGGAGGAG ATATGAGGGA
961 CAATTGGAGA AGTGAATTAT ATAAATATAA AGTAGTAAAA ATTGAACCAT TAGGAGTAGC ACCCACCAAG GCAAAGAGAA
```

|  |  |  |  |  |  |  |  |  |
| --- | --- | --- | --- | --- | --- | --- | --- | --- |
| 1041 | GAGTGGTGCA | GAGAGAAAAA | AGAGCAGTGG | GAATAGGAGC | TTGTTCCTT | GGGTCTTGG | GAGCAGCAGG | AAGCACTATG |
| 1121 | GGCGCAGCGT | CAATGACGCT | GACGGTACAG | GCCAGACAAT | TATTGTCTGG | TATAGTGCAG | CAGCAGAACA | ATTTGCTGAG |
| 1201 | GGCTATTGAG | GCGCAACAGC | ATCTGTTGCA | ACTCACAGTC | TGGGGCATCA | AGCAGCTCCA | GGCAAGAATC | CTGGCTGTGG |
| 1281 | AAAGATACCT | AAAGGATCAA | CAGCTCCTGG | GGATTGGGG | TTGCTCTGGA | AAACTCATTT | GCACCACTGC | TGTGCCTTGG |
| 1361 | AATGCTAGTT | GGAGTAATAA | ATCTCTGGAA | CAGATTGGA | ATCACACGAC | CTGGATGGAG | TGGGACAGAG | AAATTAACAA |
| 1441 | TTACACAAGC | TTAATACTACT | CCTTAATTGA | AGAATCGCAA | AACCAGCAAG | AAAAGAATGA | ACAAGAATTA | TTGGAATTAG |
| 1521 | ATAAATGGGC | AAGTTTGTGG | AATTGGTTTA | ACATAACAAA | TTGGCTGTGG | TATATAAAAT | TATTCATAAT | GATAGTAGGA |
| 1601 | GGCTTGGTAG | GTTTAAGAAT | AGTTTTTGTCT | GTACTTTCTA | TAGTGAATAG | AGTTAGGCAG | GGATATTCAC | CATTATCGTT |
| 1681 | TCAGACCCAC | CTCCCAACCC | CGAGGGGACC | CGACAGGCC | GAAGGAATAG | AAGAAGAAGG | TGGAGAGAGA | GACAGAGACA |
| 1761 | GATCCATTCG | ATTAGTGAAC | GGATCTCGAC | GGTATCGCTA | GCTTTTAAAA | GAAAAGGGGG | GATTGGGGGG | TACAGTGCAG |
| 1841 | GGGAAAGAAT | AGTAGACATA | ATAGCAACAG | ACATACAAAC | TAAAGAATTA | CAAAAACAAA | TTACAAAAT | TCAAAATTTT |
| 1921 | ACTAGTGAGG | GCCTATTTCC | CATGATTCTT | TCATATTTGC | ATATACGATA | CAAGGCTGTT | AGAGAGATAA | TTGGAATTAA |
| 2001 | TTTGAAGTGA | AACACAAAGA | TATTAGTACA | AAATACGTGA | CGTAGAAAGT | AATAATTCTT | TGGGTAGTTT | GCAGTTTTAA |
| 2081 | AATTATGTTT | TAAATAGGAC | TATCATATGC | TTACCGTAAC | TTGAAAGTAT | TTCGATTCTT | TGGCTTTATA | TATCTTGTGG |
| 2161 | AAAGGACGAA | ACACCGGTGA | ATCCTTGCCC | AGGTGCATTC | TCGAGAATGC | ACCTGGGCAA | GGATTCTATT | TTGAATTCCA |
| 2241 | ACTTTGTATA | GAAAAGTTGG | GGTTGCGCCT | TTTCCAAGGC | AGCCCTGGGT | TTGCGCAGGG | ACGCGGCTGC | TCTGGGCGTG |
| 2321 | GTTCCGGGAA | ACGCAGCGGC | GCCGACCTTG | GGTCTCGCAC | ATTCTTCACG | TCCGTTTCGA | GCGTCACCCG | GATCTTCGCC |
| 2401 | GCTACCTTGG | TGGGCCCCC | GGCGACGCTT | CCTGCTCCGC | CCCTAAGTCG | GGAAGGTTCC | TTGCGGTTTC | CGGCGTGCCG |
| 2481 | GACGTGACAA | ACGGAAGCCG | CACGTCTCAC | TAGTACCCTC | GCAGACGGAC | AGCGCCAGGG | AGCAATGGCA | GCGCGCCGAC |
| 2561 | CGCGATGGGC | TGTGGCCAAT | AGCGGCTGCT | CAGCAGGGCG | CGCCGAGAGC | AGCGGCCGGG | AAGGGGCGGT | GCGGGAGGCG |
| 2641 | GGGTGTGGGG | CGGTAGTGTG | GGCCCTGTTC | CTGCCCAGCG | GGTGTTCGCG | ATTCTGCAAG | CCTCCGAGC | GCACGTGCGC |
| 2721 | AGTCGGCTCC | CTCGTTGACC | GAATCACCGA | CCTCTCTCCC | CAGGCAAGTT | TGTACAAAAA | AGCAGGCTGC | CACCATGGTG |
| 2801 | AGCAAGGGCG | AGGAGCTGTT | CACCGGGGTG | GTGCCCATCC | TGGTCGAGCT | GGACGGCGAC | GTAAACGGCC | ACAAGTTCAG |
| 2881 | CGTGTCCGGC | GAGGGCGAGG | GCGATGCCAC | CTACGGCAAG | CTGACCCTGA | AGTTCATCTG | CACCACCGGC | AAGCTGCCCG |
| 2961 | TGCCCTGGCC | CACCCTCGTG | ACCACCCTGA | CCTACGGCGT | GCAGTGCTTC | AGCCGCTACC | CCGACCACAT | GAAGCAGCAC |
| 3041 | GACTTCTTCA | AGTCCGCCAT | GCCCGAAGGC | TACGTCCAGG | AGCGCACCAT | CTTCTTCAAG | GACGACGGCA | ACTACAAGAC |
| 3121 | CCGCGCCGAG | GTGAAGTTTC | AGGGCGACAC | CCTGGTGAAC | CGCATCGAGC | TGAAGGGCAT | CGACTTCAAG | GAGGACGGCA |
| 3201 | ACATCTTGGG | GCACAAGCTG | GAGTACAAC | ACAACAGCCA | CAACGTCTAT | ATCATGGCCG | ACAAGCAGAA | GAACGGCATC |
| 3281 | AAGGTGAAC | TCAAGATCCG | CCACAACATC | GAGGACGGCA | GCGTGCAGCT | CGCCGACCAC | TACCAGCAGA | ACACCCCAT |
| 3361 | CGGCGACGGC | CCCGTGCTGC | TGCCCGACAA | CCACTACCTG | AGCACCCAGT | CCGCCCTGAG | CAAAGACCCC | AACGAGAAGC |
| 3441 | GCGATCACAT | GGTCTTGCTG | GAGTTCGTGA | CCGCCGCCGG | GATCACTCTC | GGCATGGACG | AGCTGTACAA | GTAACCCAG |
| 3521 | CTTCTCTGTA | CAAAGTGGTG | GTACCCGATA | ATCAACCTCT | GGATTACAAA | ATTTGTGAAA | GATTGACTGG | TATTCTTAAC |
| 3601 | TATGTGCTC | CTTTTACGCT | ATGTGGATAC | GCTGCTTTAA | TGCTTTTGTA | TCATGCTATT | GCTTCCCGTA | TGGCTTTTAT |
| 3681 | TTTCTCCTCC | TTGTATAAAT | CCTGGTTGCT | GTCTCTTTAT | GAGGAGTTGT | GGCCCGTTGT | CAGGCAACGT | GGCGTGGTGT |
| 3761 | GCACTGTGTT | TGCTGACGCA | ACCCCACTG | GTGGGGGCAT | TGCCACCACC | TGTCAGCTCC | TTTCCGGGAC | TTTCGCTTTC |
| 3841 | CCCTCCCTA | TTGCCACGGC | GGAACATCAT | GCCGCTGCCC | TTGCCCGCTG | CTGGACAGGG | GCTCGGCTGT | TGGGCACTGA |
| 3921 | CAATTCCGTG | GTGTTGTTCG | GGAAGCTGAC | GTCTTTTCCA | TGGCTGCTCG | CCTGTGTTGC | CACCTGGATT | TTGCGCGGGA |
| 4001 | CGTCTTCTG | CTACGTCCCT | TCGGCCCTCA | ATCCAGCGGA | CCTTCTTCTC | CGCGGCTGCT | TGCCGCTCT | GCGGCCTCTT |
| 4081 | CCGCGTCTTC | GCCTTCGCCC | TCAGACGAGT | CGGATCTCCC | TTTGGGCCGC | CTCCCCGCAT | CGGCTTTAAG | ACCAATGACT |
| 4161 | TACAAGGCAG | CTGTAGATCT | TAGCCACTTT | TTAAAGAAAA | AGGGGGGACT | GGAAGGGCTA | ATTTACTCCC | AACGAAGACA |
| 4241 | AGATCTGCTT | TTTGCTTGTA | CTGGGTCTCT | CTGGTTAGAC | CAGATCTGAG | CCTGGGAGCT | CTCTGGCTAA | CTAGGGAACC |
| 4321 | CACGTCTTAA | GCCTCAATAA | AGCTTGCCTT | GAGTGCTTCA | AGTAGTGTGT | GCCCCGCTGT | TGTGTGACTC | TGGTAACCTAG |
| 4401 | AGATCCCTCA | GACCCCTTTA | GTCAGTGTGG | AAAATCTCTA | GCAGTAGTAG | TTCATGTCAT | CTTATTATTC | AGTATTTATA |
| 4481 | ACTTGCAAG | AAATGAATAT | CAGAGAGTGA | GAGGAACCTG | TTTATTGCGA | CTTATAATGG | TTACAAATAA | AGCAATAGCA |
| 4561 | TCACAAATTT | CACAAATAAA | GCATTTTTTT | CACTGCATTC | TAGTTGTGGT | TTGTCCAAAC | TCATCAATGT | ATCTTATCAT |
| 4641 | GTCTGGCTCT | AGCTATCCCG | CCCCTAACTC | CGCCCATCCC | GCCCCTAACT | CCGCCAGTT | CCGCCCATTC | TCCGCCCAT |
| 4721 | GGCTGACTAA | TTTTTTTTTAT | TTATGCAGAG | GCCGAGGCCG | CCTCGGCCCTC | TGAGCTATTC | CAGAAGTAGT | GAGGAGGCTT |

```
4801 TTTTGGAGGC CTAGGGACGT ACCCAATTCG CCCTATAGTG AGTCGTATTA CGCGCGCTCA CTGGCCGTCG TTTTACAACG
4881 TCGTGACTGG GAAAACCCTG GCGTTACCCA ACTTAATCGC CTTGCAGCAC ATCCCCCTTT CGCCAGCTGG CGTAATAGCG
4961 AAGAGGCCCG CACCGATCGC CCTTCCCAAC AGTTGCGCAG CCTGAATGGC GAATGGGACG CGCCCTGTAG CGGCGCATTA
5041 AGCGCGGCGG GTGTGGTGGT TACGCGCAGC GTGACCGCTA CACTTGCCAG CGCCCTAGCG CCCGCTCCTT TCGCTTTCTT
5121 CCCTTCCTTT CTCGCCACGT TCGCCGGCTT TCCCCGTCAA GCTCTAAATC GGGGGCTCCC TTTAGGGTTC CGATTTAGTG
5201 CTTTACGGCA CCTCGACCCC AAAAAACTTG ATTAGGGTGA TGGTTCACGT AGTGGGCCAT CGCCCTGATA GACGGTTTTT
5281 CGCCCTTTGA CGTTGGAGTC CACGTTCTTT AATAGTGGAC TCTTGTTCCA AACTGGAACA ACACTCAACC CTATCTCGGT
5361 CTATTCTTTT GATTTATAAG GGATTTTGCC GATTTCGGCC TATTGGTTAA AAAATGAGCT GATTTAACAA AAATTTAACG
5441 CGAATTTTAA CAAAATATTA ACGCTTACAA TTTAGGTGGC ACTTTTCGGG GAAATGTGCG CGGAACCCCT ATTTGTTTAT
5521 TTTTCTAAAT ACATTCAAAT ATGTATCCGC TCATGAGACA ATAACCCTGA TAAATGCTTC AATAATATTG AAAAAGGAAG
5601 AGTATGAGTA TTCAACATTT CCGTGTCGCC CTTATTCCCT TTTTTGCGGC ATTTTGCCTT CCTGTTTTTG CTCACCCAGA
5681 AACGCTGGTG AAAGTAAAAG ATGCTGAAGA TCAGTTGGGT GCACGAGTGG GTTACATCGA ACTGGATCTC AACAGCGGTA
5761 AGATCCTTGA GAGTTTTCGC CCCGAAGAAC GTTTTCCAAT GATGAGCACT TTTAAAGTTC TGCTATGTGG CGCGGTATTA
5841 TCCCGTATTG ACGCCGGGCA AGAGCAACTC GGTCGCCGCA TACACTATTC TCAGAATGAC TTGGTTGAGT ACTCACCAGT
5921 CACAGAAAAG CATCTTACGG ATGGCATGAC AGTAAGAGAA TTATGCAGTG CTGCCATAAC CATGAGTGAT AACACTGCGG
6001 CCAACTTACT TCTGACAACG ATCGGAGGAC CGAAGGAGCT AACCGCTTTT TTGCACAACA TGGGGGATCA TGTAACTCGC
6081 CTTGATCGTT GGGAACCGGA GCTGAATGAA GCCATACCAA ACGACGAGCG TGACACCACG ATGCCTGTAG CAATGGCAAC
6161 AACGTTGCGC AAACTATTAA CTGGCGAACT ACTTACTCTA GCTTCCCGGC AACAATTAAT AGACTGGATG GAGGCGGATA
6241 AAGTTGCAGG ACCACTTCTG CGCTCGGCCC TTCCGGCTGG CTGGTTTATT GCTGATAAAT CTGGAGCCGG TGAGCGTGGG
6321 TCTCGCGGTA TCATTGCAGC ACTGGGGCCA GATGGTAAGC CCTCCCGTAT CGTAGTTATC TACACGACGG GGAGTCAGGC
6401 AACTATGGAT GAACGAAATA GACAGATCGC TGAGATAGGT GCCTCACTGA TTAAGCATTG GTAACTGTCA GACCAAGTTT
6481 ACTCATATAT ACTTTAGATT GATTTAAAAC TTCATTTTTA ATTTAAAAGG ATCTAGGTGA AGATCCTTTT TGATAATCTC
6561 ATGACCAAAA TCCCTTAACG TGAGTTTTCG TTCCACTGAG CGTCAGACCC CGTAGAAAAG ATCAAAGGAT CTTCTTGAGA
6641 TCCTTTTTTT CTGCGCGTAA TCTGCTGCTT GCAAACAAAA AAACCACCGC TACCAGCGGT GGTTTGTTTG CCGGATCAAG
6721 AGCTACCAAC TCTTTTTCCG AAGGTAACTG GCTTCAGCAG AGCGCAGATA CCAAATACTG TTCTTCTAGT GTAGCCGTAG
6801 TTAGGCCACC ACTTCAAGAA CTCTGTAGCA CCGCCTACAT ACCTCGCTCT GCTAATCCTG TTACCAGTGG CTGCTGCCAG
6881 TGGCGATAAG TCGTGTCTTA CCGGGTTGGA CTCAAGACGA TAGTTACCGG ATAAGGCGCA GCGGTCGGGC TGAACGGGGG
6961 GTTCGTGCAC ACAGCCCAGC TTGGAGCGAA CGACCTACAC CGAACTGAGA TACCTACAGC GTGAGCTATG AGAAAGCGCC
7041 ACGCTTCCCG AAGAGAGAAA GGCGGACAGG TATCCGGTAA GCGGCAGGGT CGGAACAGGA GAGCGCACGA GGGAGCTTCC
7121 AGGGGGAAAC GCCTGGTATC TTTATAGTCC TGTCGGGTTT CGCCACCTCT GACTTGAGCG TCGATTTTTG TGATGCTCGT
7201 CAGGGGGGCG GAGCCTATGG AAAAACGCCA GCAACGCGGC CTTTTTACGG TTCCTGGCCT TTTGCTGGCC TTTTGCTCAC
7281 ATGTTCTTTC CTGCGTTATC CCCTGATTCT GTGGATAACC GTATTACCGC CTTTGAGTGA GCTGATACCG CTCGCCGCAG
7361 CCGAACGACC GAGCGCAGCG AGTCAGTGAG CGAGGAAGCG GAAGAGCGCC CAATACGCAA ACCGCCTCTC CCCGCGCGTT
7441 GGCCGATTCA TTAATGCAGC TGGCACGACA GGTTTCCCGA CTGGAAAGCG GGCAGTGAGC GCAACGCAAT TAATGTGAGT
7521 TAGCTCACTC ATTAGGCACC CCAGGCTTTA CACTTTATGC TTCCGGCTCG TATGTTGTGT GGAATTGTGA GCGGATAACA
7601 ATTTCACACA GGAAACAGCT ATGACCATGA TTACGCCAAG CGCGCAATTA ACCCTCACTA AAGGGAACAA AAGCTGGAGC
7681 TGCAAGCTT
```

### Validation by Restriction Enzyme Digestion

| Restriction Enzymes | Cutting Sites | DNA Fragments (bp) |
| --- | --- | --- |
| XhoI | 2201 | 7689 |
| ApaLI | 3760, 5720, 6966 | 1960, 1246, 4483 |
| ApaLI+XhoI | 2201, 3760, 5720, 6966 | 1559, 1960, 1246, 2924 |

### Vector Summary

|  |  |
| --- | --- |
| Vector ID | VB231004-1661ewa |
| Vector Name | pLV[Exp]-SYN1>EGFP:{miRNA-502-3p sponge} |
| Vector Size | 7602 bp |
| Viral Genome Size | 4127 bp |
| Vector Type | Mammalian Gene Expression Lentiviral Vector |
| Plasmid Copy Number | High |
| Antibiotic Resistance | Ampicillin |
| Cloning Host | VB UltraStable (or alternative strain) |

### Vector Map

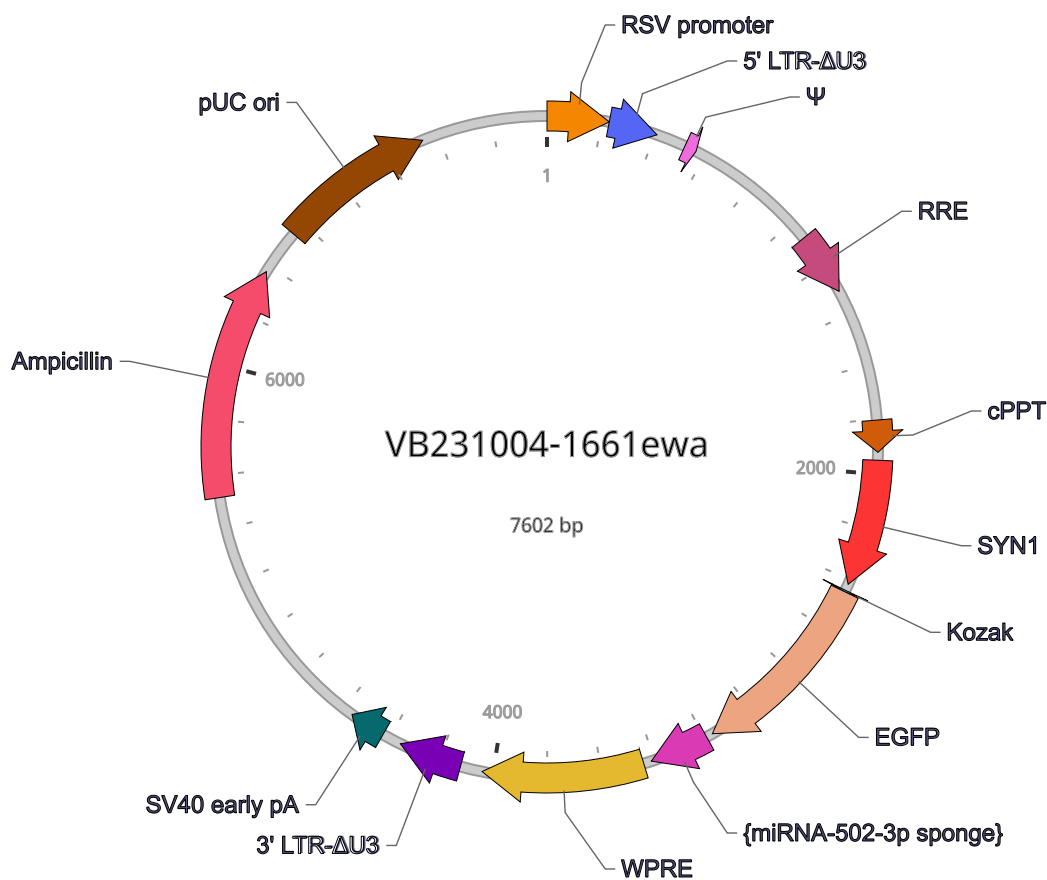

### Vector Components

| Name | Position | Size (bp) | Type | Description | Application notes |
| --- | --- | --- | --- | --- | --- |
| RSV promoter | 1-229 | 229 | Promoter | *None* | Strong promoter; drives transcription of viral RNA in packaging cells. |
| 5' LTR-ΔU3 | 230-410 | 181 | LTR | *None* | Allows transcription of viral RNA and its packaging into virus. |
| Ψ | 521-565 | 45 | misc_feature | *None* | Allows packaging of viral RNA into virus. |
| RRE | 1075-1308 | 234 | misc_feature | *None* | Rev protein binding site that allows Rev-dependent nuclear export of viral RNA during viral packaging. |
| cPPT | 1803-1920 | 118 | misc_feature | *None* | Facilitates the nuclear import of HIV-1 cDNA through a central DNA flap. |
| SYN1 | 1950-2418 | 469 | Promoter | *None* | Tissue specificity: Brain. Cell type specificity: Mature neurons. |
| Kozak | 2443-2448 | 6 | misc_feature | *None* | Facilitates translation initiation of ATG start codon downstream of the Kozak sequence. |
| EGFP | 2449-3168 | 720 | CDS | *None* | Commonly used green fluorescent protein; ranked high in brightness, photostability and pH stability among all fluorescent proteins. |
| {miRNA-502-3p sponge} | 3193-3413 | 221 | CDS | *None* | *None* |
| WPRE | 3443-4040 | 598 | misc_feature | *None* | Enhances virus stability in packaging cells, leading to higher titer of packaged virus; enhances higher expression of transgenes. |
| 3' LTR-ΔU3 | ■ 4122-4356 | 235 | LTR | *None* | Allows packaging of viral RNA into virus; self-inactivates the 5' LTR by a copying mechanism during viral genome integration; contains polyadenylation signal for transcription termination. |
| SV40 early pA | ■ 4429-4563 | 135 | polyA_signal | *None* | Allows transcription termination and polyadenylation of mRNA transcribed by Pol II RNA polymerase. |
| Ampicillin | ■ 5517-6377 | 861 | CDS | *None* | Allows E. coli to be resistant to ampicillin. |
| pUC ori | ■ 6548-7136 | 589 | rep_origin | *None* | Facilitates plasmid replication in E. coli; regulates high-copy plasmid number (500-700). |

Note: Components added by user are listed in **bold red** text.

### Vector Sequence

```
1  AATGTAGTCT TATGCAATAC TCTTGTAGTC TTGCAACATG GTAACGATGA GTTAGCAACA TGCCTTACAA GGAGAGAAAA
81  AGCACCGTGC ATGCCGATTG GTGGAAGTAA GGTGGTACGA TCGTGCCTTA TTAGGAAGGC AACAGACGGG TCTGACATGG
161 ATTGGACGAA CCACTGAATT GCCGCATTGC AGAGATATTG TATTTAAGTG CCTAGCTCGA TACATAAACG GGTCTCTCTG
241 GTTAGACCAG ATCTGAGCCT GGGAGCTCTC TGGCTAACTA GGGAACCCAC TGCTTAAGCC TCAATAAAGC TTGCCTTGAG
321 TGCTTCAAGT AGTGTGTGCC CGTCTGTTGT GTGACTCTGG TAACTAGAGA TCCCTCAGAC CCTTTTAGTC AGTGTGGAAA
401 ATCTCTAGCA GTGGCGCCCG AACAGGGACT TGAAAGCGAA AGGGAAACCA GAGGAGCTCT CTCGACGCAG GACTCGGCTT
481 GCTGAAGCGC GCACGGCAAG AGGCGAGGGG CGGCGACTGG TGAGTACGCC AAAAATTTTG ACTAGCGGAG GCTAGAAGGA
561 GAGAGATGGG TGCGAGAGCG TCAGTATTAA GCGGGGGAGA ATTAGATCGC GATGGGAAAA AATTCGGTTA AGGCCAGGGG
641 GAAAGAAAAA ATATAAATTA AAACATATAG TATGGGCAAG CAGGGAGCTA GAACGATTCG CAGTTAATCC TGGCCTGTTA
721 GAAACATCAG AAGGCTGTAG ACAAATACTG GGACAGCTAC AACCATCCCT TCAGACAGGA TCAGAAGAAC TTAGATCATT
801 ATATAATACA GTAGCAACCC TCTATTGTGT GCATCAAAGG ATAGAGATAA AAGACACCAA GGAAGCTTTA GACAAGATAG
881 AGGAAGAGCA AAACAAAAGT AAGACCACCG CACAGCAAGC GGCCGCTGAT CTTCAGACCT GGAGGAGGAG ATATGAGGGA
961 CAATTGGAGA AGTGAATTAT ATAAATATAA AGTAGTAAAA ATTGAACCAT TAGGAGTAGC ACCCACCAAG GCAAAGAGAA
1041 GAGTGGTGCA GAGAGAAAAA AGAGCAGTGG GAATAGGAGC TTTGTTCCTT GGGTTCTTGG GAGCAGCAGG AAGCACTATG
1121 GGCGCAGCGT CAATGACGCT GACGGTACAG GCCAGACAAT TATTGTCTGG TATAGTGCAG CAGCAGAACA ATTTGCTGAG
1201 GGCTATTGAG GCGCAACAGC ATCTGTTGCA ACTCACAGTC TGGGGCATCA AGCAGCTCCA GGCAAGAATC CTGGCTGTGG
1281 AAAGATACCT AAAGGATCAA CAGCTCCTGG GGATTTGGGG TTGCTCTGGA AAACTCATTT GCACCACTGC TGTGCCTTGG
1361 AATGCTAGTT GGAGTAATAA ATCTCTGGAA CAGATTTGGA ATCACACGAC CTGGATGGAG TGGGACAGAG AAATTAACAA
```

```
1441 TTACACAAGC TTAATACACT CCTTAATTGA AGAATCGCAA AACCAGCAAG AAAAGAATGA ACAAGAATTA TTGGAATTAG
1521 ATAAATGGGC AAGTTTGTGG AATTGGTTTA ACATAACAAA TTGGCTGTGG TATATAAAAT TATTCATAAT GATAGTAGGA
1601 GGCTTGGTAG GTTTAAGAAT AGTTTTTGCT GTACTTTCTA TAGTGAATAG AGTTAGGCAG GGATATTCAC CATTATCGTT
1681 TCAGACCCAC CTCCCAACCC CGAGGGGACC CGACAGGCCC GAAGGAATAG AAGAAGAAGG TGGAGAGAGA GACAGAGACA
1761 GATCCATTCG ATTAGTGAAC GGATCTCGAC GGTATCGCTA GCTTTTAAAA GAAAAGGGGG GATTGGGGGG TACAGTGCAG
1841 GGGAAAGAAT AGTAGACATA ATAGCAACAG ACATACAAAC TAAAGAATTA CAAAAACAAA TTACAAAAAT TCAAAATTTT
1921 ACTAGTATCA ACTTTGTATA GAAAAGTTGC TGCAGAGGGC CCTGCGTATG AGTGCAAGTG GGTTTTAGGA CCAGGATGAG
2001 GCGGGGTGGG GGTGCCTACC TGACGACCGA CCCCGACCCA CTGGACAAGC ACCCAACCCC CATTCCCCAA ATTGCGCATC
2081 CCCTATCAGA GAGGGGGAGG GGAAACAGGA TGCGGCGAGG CGCGTGCGCA CTGCCAGCTT CAGCACCGCG GACAGTGCCT
2161 TCGCCCCCGC CTGGCGGCGC GCGCCACCGC CGCCTCAGCA CTGAAGGCGC GCTGACGTCA CTCGCCGGTC CCCCGCAAAC
2241 TCCCCTTCCC GGCCACCTTG GTCGCGTCCG CGCCGCCGCC GGCCCAGCCG GACCGCACCA CGCGAGGCGC GAGATAGGGG
2321 GGCACGGGCG CGACCATCTG CGCTGCGGCG CCGGCGACTC AGCGCTGCCT CAGTCTGCGG TGGGCAGCGG AGGAGTCGTG
2401 TCGTGCCTGA GAGCGCAGCA AGTTTGTACA AAAAAGCAGG CTGCCACCAT GGTGAGCAAG GGCGAGGAGC TGTTCACCGG
2481 GGTGGTGCCC ATCCTGGTCG AGCTGGACGG CGACGTAAAC GGCCACAAGT TCAGCGTGTC CGGCGAGGGC GAGGGCGATG
2561 CCACCTACGG CAAGCTGACC CTGAAGTTCA TCTGCACCAC CGGCAAGCTG CCCGTGCCCT GGCCCACCCT CGTGACCACC
2641 CTGACCTACG GCGTGCAGTG CTTCAGCCGC TACCCCGACC ACATGAAGCA GCACGACTTC TTCAAGTCCG CCATGCCCGA
2721 AGGCTACGTC CAGGAGCGCA CCATCTTCTT CAAGGACGAC GGCAACTACA AGACCCGCGC CGAGGTGAAG TTCGAGGGCG
2801 ACACCCTGGT GAACCGCATC GAGCTGAAGG GCATCGACTT CAAGGAGGAC GGCAACATCC TGGGGCACAA GCTGGAGTAC
2881 AACTACAACA GCCACAACGT CTATATCATG GCCGACAAGC AGAAGAACGG CATCAAGGTG AACTTCAAGA TCCGCCACAA
2961 CATCGAGGAC GGCAGCGTGC AGCTCGCCGA CCACTACCAG CAGAACACCC CCATCGGCGA CGGCCCCGTG CTGCTGCCCG
3041 ACAACCACTA CCTGAGCACC CAGTCCGCCC TGAGCAAAGA CCCCAACGAG AAGCGCGATC ACATGGTCCT GCTGGAGTTC
3121 GTGACCGCCG CCGGGATCAC TCTCGGCATG GACGAGCTGT ACAAGTAAAC CCAGCTTTCT TGTACAAAGT GGCTCGAGCC
3201 GGTGAATCCT TGGGTGGTGC ATTCCGGTGA ATCCTTGGGT GGTGCATTCC GGTGAATCCT TGGGTGGTGC ATTCCGGTGA
3281 ATCCTTGGGT GGTGCATTCC GGTGAATCCT TGGGTGGTGC ATTCCGGTGA ATCCTTGGGT GGTGCATTCC GGTGAATCCT
3361 TGGGTGGTGC ATTCCGGTGA ATCCTTGGGT GGTGCATTCC GGATCGCGGG CCCCAACTTT ATTATACATA GTTGATCAAT
3441 TCCGATAATC AACCTCTGGA TTACAAAATT TGTGAAAGAT TGACTGGTAT TCTTAACTAT GTTGCTCCTT TTACGCTATG
3521 TGGATACGCT GCTTTAATGC CTTTGTATCA TGCTATTGCT TCCCGTATGG CTTTCATTTT CTCCTCCTTG TATAAATCCT
3601 GGTTGCTGTC TCTTTATGAG GAGTTGTGGC CCGTTGTCAG GCAACGTGGC GTGGTGTGCA CTGTGTTTGC TGACGCAACC
3681 CCCACTGGTT GGGGCATTGC CACCACCTGT CAGCTCCTTT CCGGGACTTT CGCTTTCCCC CTCCCTATTG CCACGGCGGA
3761 ACTCATCGCC GCCTGCCTTG CCCGCTGCTG GACAGGGGCT CGGCTGTTGG GCACTGACAA TTCCGTGGTG TTGTCGGGGA
3841 AGCTGACGTC CTTTCCATGG CTGCTCGCCT GTGTTGCCAC CTGGATTCTG CGCGGGACGT CCTTCTGCTA CGTCCCTTCG
3921 GCCCTCAATC CAGCGGACCT TCCTTCCCGC GGCCTGCTGC CGGCTCTGCG GCCTCTTCCG CGTCTTCGCC TTCGCCCTCA
4001 GACGAGTCGG ATCTCCCTTT GGGCCGCCTC CCCGCATCGG GAATTCCCGC GGTTCGCTTT AAGACCAATG ACTTACAAGG
4081 CAGCTGTAGA TCTTAGCCAC TTTTTAAAAG AAAAGGGGGG ACTGGAAGGG CTAATTCACT CCCAACGAAG ACAAGATCTG
4161 CTTTTTGCTT GTACTGGGTC TCTCTGGTTA GACCAGATCT GAGCCTGGGA GCTCTCTGGC TAACTAGGGA ACCCACTGCT
4241 TAAGCCTCAA TAAAGCTTGC CTTGAGTGCT TCAAGTAGTG TGTGCCCGTC TGTTGTGTGA CTCTGGTAAC TAGAGATCCC
4321 TCAGACCCTT TTAGTCAGTG TGGAAAATCT CTAGCAGTAG TAGTTCATGT CATCTTATTA TTCAGTATTT ATAACTTGCA
4401 AAGAAATGAA TATCAGAGAG TGAGAGGAAC TTGTTTATTG CAGCTTATAA TGGTTACAAA TAAAGCAATA GCATCACAAA
4481 TTTCACAAAT AAAGCATTTT TTTCACTGCA TTCTAGTTGT GGTTTGTCCA AACTCATCAA TGTATCTTAT CATGTCTGGC
4561 TCTAGCTATC CCGCCCCTAA CTCCGCCCAT CCCGCCCCTA ACTCCGCCCA GTTCCGCCCA TTCTCCGCCC CATGGCTGAC
4641 TAATTTTTTT TATTTATGCA GAGGCCGAGG CCGCCTCGGC CTCTGAGCTA TTCCAGAAGT AGTGAGGAGG CTTTTTTGGA
4721 GGCCTAGGGA CGTACCCAAT TCGCCCTATA GTGAGTCGTA TTACGCGCGC TCACTGGCCG TCGTTTTACA ACGTCGTGAC
4801 TGGGAAAACC CTGGCGTTAC CCAACTTAAT CGCCTTGCAG CACATCCCCC TTTCGCCAGC TGGCGTAATA GCGAAGAGGC
4881 CCGCACCGAT CGCCCTTCCC AACAGTTGCG CAGCCTGAAT GGCGAATGGG ACGCGCCCTG TAGCGGCGCA TTAAGCGCGG
4961 CGGGTGTGGT GGTTACGCGC AGCGTGACCG CTACACTTGC CAGCGCCCTA GCGCCCGCTC CTTTCGCTTT CTTCCCTTCC
5041 TTTCTCGCCA CGTTCGCCGG CTTTCCCCGT CAAGCTCTAA ATCGGGGGCT CCCTTTAGGG TTCCGATTTA GTGCTTTACG
5121 GCACCTCGAC CCAAAAAAAC TTGATTAGGG TGATGGTTCA CGTAGTGGG CATCGCCCTG ATAGACGGTT TTTGCCCCCT
5201 TGACGTTGGA GTCCACGTTC TTTAATAGTG GACTCTTGTT CCAAACTGGA ACAACACTCA ACCCTATCTC GGTCTATTCT
```

```
5281 TTTGATTTAT AAGGGATTTT GCCGATTTCG GCCTATTGGT TAAAAAATGA GCTGATTTAA CAAAAATTTA ACGCGAATTT
5361 TAACAAAATA TTAACGCTTA CAATTTAGGT GGCACTTTTC GGGGAAATGT GCGCGGAACC CCTATTTGTT TATTTTTCTA
5441 AATACATTCA AATATGTATC CGCTCATGAG ACAATAACCC TGATAAATGC TTCAATAATA TTGAAAAAGG AAGAGTATGA
5521 GTATTCAACA TTTCCGTGTC GCCCTTATTC CCTTTTTTGC GGCATTTTGC CTTCCTGTTT TTGCTCACCC AGAAACGCTG
5601 GTGAAAGTAA AAGATGCTGA AGATCAGTTG GGTGCACGAG TGGGTTACAT CGAACTGGAT CTCAACAGCG GTAAGATCCT
5681 TGAGAGTTTT CGCCCCGAAG AACGTTTTCC AATGATGAGC ACTTTTAAAG TTCTGCTATG TGGCGCGGTA TTATCCCGTA
5761 TTGACGCCGG GCAAGAGCAA CTCGGTCGCC GCATACACTA TTCTCAGAAT GACTTGGTTG AGTACTCACC AGTCACAGAA
5841 AAGCATCTTA CGGATGGCAT GACAGTAAGA GAATTATGCA GTGCTGCCAT AACCATGAGT GATAACACTG CGGCCAACTT
5921 ACTTCTGACA ACGATCGGAG GACCGAAGGA GCTAACCGCT TTTTTGCACA ACATGGGGGA TCATGTAACT CGCCTTGATC
6001 GTTGGGAACC GGAGCTGAAT GAAGCCATAC CAAACGACGA GCGTGACACC ACGATGCCTG TAGCAATGGC AACAACGTTG
6081 CGCAAACTAT TAACTGGCGA ACTACTTACT CTAGCTTCCC GGCAACAATT AATAGACTGG ATGGAGGCGG ATAAAGTTGC
6161 AGGACCACTT CTGCGCTCGG CCCTTCCGGC TGGCTGGTTT ATTGCTGATA AATCTGGAGC CGGTGAGCGT GGGTCTCGCG
6241 GTATCATTGC AGCACTGGGG CCAGATGGTA AGCCCTCCCG TATCGTAGTT ATCTACACGA CGGGGAGTCA GGCAACTATG
6321 GATGAACGAA ATAGACAGAT CGCTGAGATA GGTGCCTCAC TGATTAAGCA TTGGTAACTG TCAGACCAAG TTTACTCATA
6401 TATACTTTAG ATTGATTTAA AACTTCATTT TTAATTTAAA AGGATCTAGG TGAAGATCCT TTTTGATAAT CTCATGACCA
6481 AAATCCCTTA ACGTGAGTTT TCGTTCCACT GAGCGTCAGA CCCCGTAGAA AAGATCAAAG GATCTTCTTG AGATCCTTTT
6561 TTTCTGCGCG TAATCTGCTG CTTGCAAACA AAAAAACCAC CGCTACCAGC GGTGGTTTGT TTGCCGGATC AAGAGCTACC
6641 AACTCTTTTT CCGAAGGTAA CTGGCTTCAG CAGAGCGCAG ATACCAAATA CTGTTCTTCT AGTGTAGCCG TAGTTAGGCC
6721 ACCACTTCAA GAACTCTGTA GCACCGCCTA CATACCTCGC TCTGCTAATC CTGTTACCAG TGGCTGCTGC CAGTGGCGAT
6801 AAGTCGTGTC TTACCGGGTT GGACTCAAGA CGATAGTTAC CGGATAAGGC GCAGCGGTCG GGCTGAACGG GGGGTTCGTG
6881 CACACAGCCC AGCTTGGAGC GAACGACCTA CACCGAACTG AGATACCTAC AGCGTGAGCT ATGAGAAAGC GCCACGCTTC
6961 CCGAAGAGAG AAAGGCGGAC AGGTATCCGG TAAGCGGCAG GGTCGGAACA GGAGAGCGCA CGAGGGAGCT TCCAGGGGGA
7041 AACGCCTGGT ATCTTTATAG TCCTGTCGGG TTTCGCCACC TCTGACTTGA GCGTCGATTT TTGTGATGCT CGTCAGGGGG
7121 GCGGAGCCTA TGGAAAAACG CCAGCAACGC GGCCTTTTTA CGGTTCCTGG CCTTTTGCTG GCCTTTTGCT CACATGTTCT
7201 TTCCTGCGTT ATCCCCTGAT TCTGTGGATA ACCGTATTAC CGCCTTTGAG TGAGCTGATA CCGCTCGCCG CAGCCGAACG
7281 ACCGAGCGCA GCGAGTCAGT GAGCGAGGAA GCGGAAGAGC GCCCAATACG CAAACCGCCT CTCCCCGCGC GTTGGCCGAT
7361 TCATTAATGC AGCTGGCACG ACAGGTTTCC CGACTGGAAA GCGGGCAGTG AGCGCAACGC AATTAATGTG AGTTAGCTCA
7441 CTCATTAGGC ACCCCAGGCT TTACACTTTA TGCTTCCGGC TCGTATGTTG TGTGGAATTG TGAGCGGATA ACAATTTCAC
7521 ACAGGAAACA GCTATGACCA TGATTACGCC AAGCGCGCAA TTAACCCTCA CTAAAGGGAA CAAAAGCTGG AGCTGCAAGC
7601 TT
```

### Validation by Restriction Enzyme Digestion

| Restriction Enzymes | Cutting Sites | DNA Fragments (bp) |
| --- | --- | --- |
| XhoI | 3194 | 7602 |
| SpeI | 1922 | 7602 |
| AvaI | 1700, 3194 | 1494, 6108 |
| ApaLI | 3657, 5633, 6879 | 1976, 1246, 4380 |
| AflII | 294, 4240 | 3946, 3656 |
| ApaLI+XhoI | 3194, 3657, 5633, 6879 | 463, 1976, 1246, 3917 |
| ApaLI+AvaI | 1700, 3194, 3657, 5633, 6879 | 1494, 463, 1976, 1246, 2423 |
| ApaLI+SpeI | 1922, 3657, 5633, 6879 | 1735, 1976, 1246, 2645 |
| ApaLI+AflII | 294, 3657, 4240, 5633, 6879 | 3363, 583, 1393, 1246, 1017 |
