## Supplementary information file 2 for "Modulation of microRNA-502-3p significantly influences synaptic activity, dendritic spine density and mitochondrial morphology in the mice brain"

**Table 1. Oligonucleotide sequences of primers used for miR-502-3p quantitative reverse transcription-polymerase chain reaction analysis**

| **Gene(s)** | **Sequence(s)** |
| --- | --- |
| U6 SnRNA | F 5ʹ CGCTTCGGCAGCACATATACTAA 3ʹ |
|  | R 5ʹ TATGGAACGCTTCACGAATTTGC 3ʹ |
| miR-502-3p | F 5ʹAATGCACCTGGGCAAGGATTCA 3ʹ |

**Table 2. Summary of antibody dilutions and conditions used in the immunoblotting analysis**

| **Marker(s)** | **Primary Antibody and Dilution(s)**  **(4^°^C, overnight)** | **Purchased from Company, City & State** | **Secondary Antibody, Dilution(s)**  **(Room temperature, 2 h)** | **Purchased from Company, City & State** |
| --- | --- | --- | --- | --- |
| Beta Actin  (66009-1-Ig) | Mouse monoclonal  1:500 | Proteintech, Rosemont, IL | Rabbit anti-mouse IgG HRP 1:10,000  (A9044-2 mL) | Millipore Sigma  Burlington, MA |
| GABARα1  (12420-1-AP) | Rabbit  polyclonal  1:500 | Proteintech, Rosemont, IL | Goat anti-rabbit IgG HRP 1:10,000  (A9169-2 mL) | Millipore Sigma  Burlington, MA |
| Gephyrin  (12681-1-AP) | Rabbit  polyclonal  1:3000 | Proteintech, Rosemont, IL | Goat anti-rabbit IgG HRP 1:10,000  (A9169-2 mL) | Millipore Sigma  Burlington, MA |
| PSD95-Specific DLG4  (20665-1-AP) | Rabbit  polyclonal | Proteintech, Rosemont, IL | Goat anti-rabbit IgG HRP 1:10,000  (A9169-2 mL) | Millipore Sigma  Burlington, MA |
| Synaptophysin  (17785-1-AP) | Rabbit  polyclonal  1:500 | Proteintech, Rosemont, IL | Rabbit anti-mouse IgG HRP 1:10,000  (A9044-2 mL) | Millipore Sigma  Burlington, MA |
